# Spatiotemporal Analysis of Lung Immune Dynamics in Lethal *Coccidioides posadasii* Infection

**DOI:** 10.1101/2024.08.21.609002

**Authors:** Oscar A. Davalos, Aimy Sebastian, Nicole F. Leon, Margarita V. Rangel, Nadia Miranda, Deepa K. Murugesh, Ashlee M. Phillips, Katrina K. Hoyer, Nicholas R. Hum, Gabriela G. Loots, Dina R. Weilhammer

## Abstract

Coccidioidomycosis, or Valley Fever, is a lung disease caused by inhalation of *Coccidioides* fungi, prevalent in the Southwestern U.S., Mexico, and parts of Central and South America. 350,000 cases are reported annually in the U.S., although that number is expected to increase as climate change expands fungal geographic range. While 60% of infections are asymptomatic, the symptomatic 40% are often misdiagnosed due to similarities with bronchitis or pneumonia. A small subset of infection progress to severe illness, necessitating a better understanding of immune responses during lethal infection. Using single-cell RNA sequencing and spatial transcriptomics, we characterized lung responses during *Coccidioides* infection. We identified monocyte-derived *Spp1*-expressing macrophages as potential mediators of tissue remodeling and fibrosis, marked by high expression of profibrotic and proinflammatory transcripts. These macrophages showed elevated TGF-β and IL-6 signaling, pathways involved in fibrosis pathogenesis. Additionally, we observed significant neutrophil infiltration and defective lymphocyte responses, indicating severe adaptive immunity dysregulation in lethal, acute infection. These findings enhance our understanding of *Coccidioides* infection and suggest new therapeutic targets.

**Importance:** Coccidioidomycosis, commonly known as Valley Fever, is a lung disease caused by the inhalation of *Coccidioides* fungi, which is prevalent in the Southwestern U.S., Mexico, and parts of Central and South America. With climate change potentially expanding the geographic range of this fungus, understanding the immune responses during severe infections is crucial. Our study used advanced techniques to analyze lung responses during *Coccidioides* infection, identifying specific immune cells that may contribute to tissue damage and fibrosis. These findings provide new insights into the disease mechanisms and suggest potential targets for therapeutic intervention, which could improve outcomes for patients suffering from severe Valley Fever.

## Introduction

Coccidioidomycosis, also known as Valley fever, is a pulmonary infectious disease caused by inhalation of soil dwelling fungi *Coccidioidies immitis* or *Coccidioides posadasii* (1–3). Coccidioidomycosis is endemic to certain regions in the Western Hemisphere including the Southwestern United States, particularly Arizona and California, as well as Mexico and parts of Central and South America (1–3). Climate change and environmental factors, such as arid soils, often influence the lifestyle of *Coccidioides* and infection rates (3). Annually, there are ∼350,0000 reported cases of Coccidioides infection in the United States, however, this number is underrepresented due to misdiagnosis, and expected to increase as climate change expands the conditions under which this fungus thrives (4). As the fungal geographic range expands and the incidence of infection rises, this once regionally confined disease is becoming a significant and widespread public health challenge (5, 6).

Roughly 60% of *Coccidioides* infections are asymptomatic, resulting in low reporting of true infection rates as diagnostic testing is not always performed. Most cases resolve spontaneously (2, 7); however, the remaining 40% of patients who do go on to become symptomatic can develop prolonged symptoms that resemble bronchitis or pneumonia. This complicates diagnosis, as these symptoms are virtually indistinguishable from those of bacterial community-acquired pneumonia (7). Unfortunately, a small subset of patients goes on to develop long lasting and potentially life threatening illnesses (8–10), highlighting the need for a better understanding of immune responses to *Coccidioides* infections. In particular, the cell types and mechanisms at play that underscore severe *Coccidioides* infection are under described.

The critical protective role of T cells in *Coccidioides* infections has been described, with Th1 and Th17 responses being essential for controlling the infection and promoting fungal clearance (11–16). In mice, vaccination with a live, attenuated Δ*cts2*/Δ*ard1*/Δ*cts3* mutant of *C. posadasii* induces effective T-cell mediated immunity, primarily through a Th17 effector response (13, 14). Similarly, another study found that vaccination with the avirulent *C. posadasii* strain, *Δcps1*, promotes long-term survival and immunity (17). In contrast, unvaccinated mice in the same study succumbed to infection. The underlying mechanisms that govern maladaptive immune responses during severe infection remain unclear in both humans and mice. However, regulatory T cells (Tregs) are associated with chronic infections likely by suppressing effective immune responses and allowing fungal persistence (18). Additionally, neutrophil numbers are notably high during chronic infections, suggesting their involvement in the ongoing inflammatory response (13, 18). While it is clear that a strong adaptive response is required to overcome infection, little is known about the mechanisms that result in insufficient protective responses during severe infection, including the effects of cellular cross talk among myeloid populations (such as resident macrophages and infiltrating monocytes) and nonimmune cells.

Utilizing two advanced technologies: single-cell RNA sequencing (scRNAseq) and spatial transcriptomics we present a longitudinal description of immune responses in lethal *Coccidioides* infection. These cutting-edge techniques provide a higher resolution understanding of lung dysregulation, allowing us to explore how the cellular composition of lung tissue is altered over time in response to infection. This work demonstrates the diverse cell type responses within the lung and suggests previously unknown roles for multiple immune cell types that may promote pathogenesis. Specifically, we identified two prominent myeloid populations, Spp1^+^ macrophages and PD-L1^+^ neutrophils, induced by infection which increase in size as the disease progresses and develop a profibrotic and proinflammatory functionality. Through this temporal characterization of pulmonary *Coccidioides* infection, we identify new potential therapeutic targets by highlighting cellular dynamics during lethal disease progression.

## Results

### *C. posadasii* infection triggers massive immune infiltration in the lung

To elucidate the cellular response in the lung during *C. posadasii* infection over time, we utilized an intranasal mouse model of infection coupled with scRNAseq. Lungs were collected from infected mice (n=5) at 5, 9, and 14 days post infection (dpi), and from uninfected control mice. Fungal burden was detectable at 5 dpi, remained steady at 9 dpi and increased by more than 100-fold by 14 dpi (Fig. 1A). Body weight dropped slightly at 9 dpi and decreased further by ∼30% of starting weight at 14 dpi, with 3 out of the 5 mice approaching humane end point for euthanasia (<70% original body weight) (Fig. 1A). Total lung cellularity increased significantly at 14 dpi, as did the relative proportion of immune cells within the lung (Fig. 1A). Histological examination of infected lungs was performed at 14 dpi, revealing increased cellularity and large clusters of immune infiltration consistent with granuloma formation (Fig. 1B-E and Fig. S1A). Our results closely recapitulated previously characterized disease progression and pathological characteristics in this model of *Coccidioides* infection (12).

**Figure 1:**
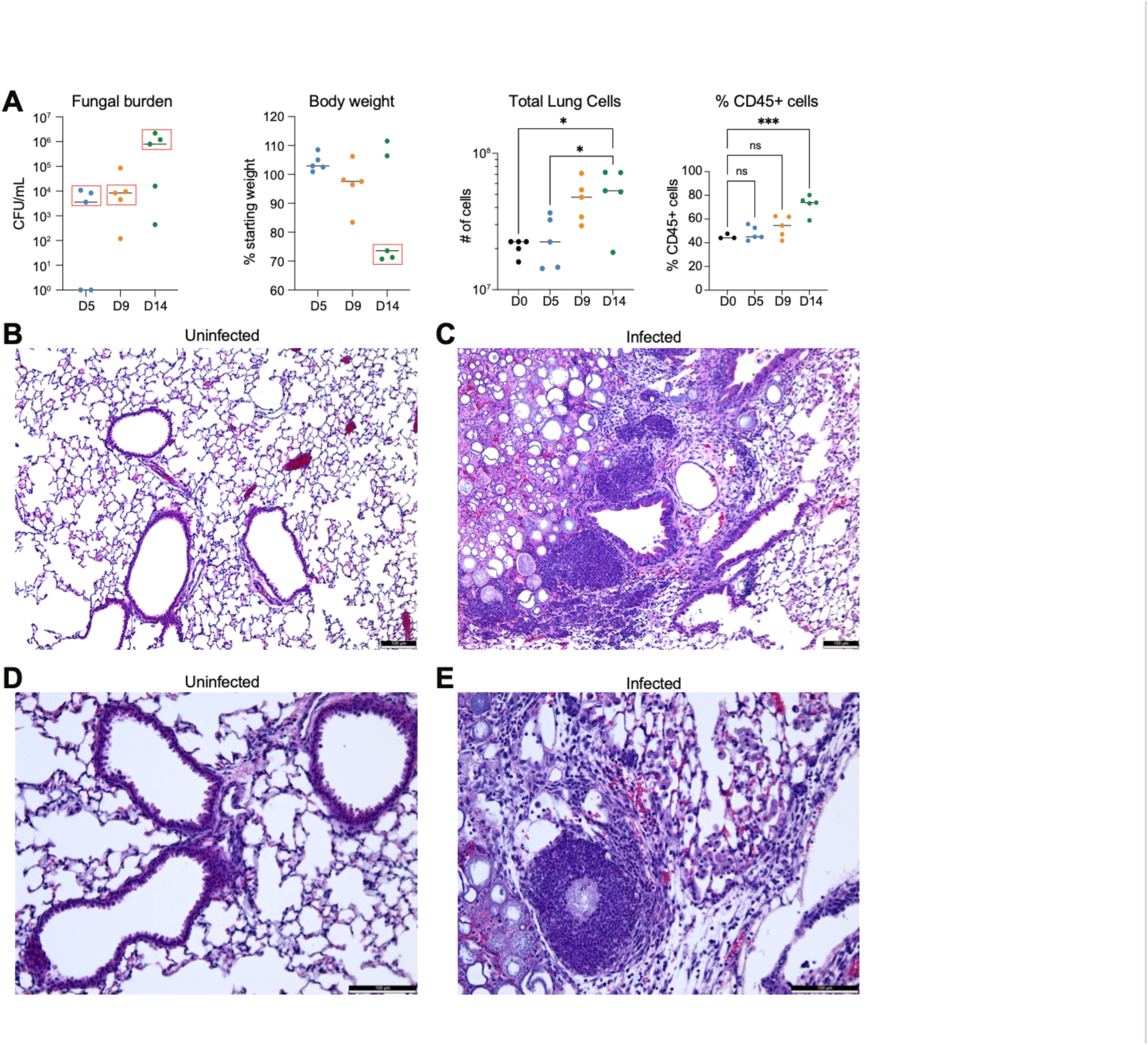
Disease progression over time is associated with increased fungal burden, increased lung cellularity and immune infiltration. (A) Lungs were harvested (n=5) at days 5, 9, and 14 post-infection (dpi) to determine the fungal burden. Red boxes indicate the animals selected for scRNAseq analysis. Body weight was measured at the time of harvest. Total lung cell counts and the percentage of CD45^+^ cells were determined by flow cytometry. (B) H&E staining of an uninfected (B and D) and infected (C and E) lung at 10x (B and C) or 20x (D and E) magnification. The white scale bar in the bottom right-hand corner represents 100 µm. *p>0.05, ***p>0.001

Next, we performed a thorough transcriptomic analysis on the infected lungs (n=3, selected for sequencing based on comparable fungal burden) using scRNAseq (Fig. 2 A). Post-filtering, we retained a total of 49,148 cells across all time points. In the lung, we identified resident and infiltrating cells, categorized into nine major cell clusters: endothelial, neutrophils, monocytes and macrophages (Mono/Mac), fibroblasts, B cells, T and natural killer cells (T/NK), epithelial, proliferating (Prolif), and mesenchymal (Fig. 2B; Fig. S1 E; and Table S1). Resident lung cells such as endothelial, fibroblasts, epithelial, and mesenchymal cells constituted 35.2%, 14.9%, 10.8%, and 2.8% of the total cell population at 0 dpi, respectively. These populations showed a decrease over time, dropping to 4.7%, 4.8%, 1.3%, and 0.16% at 14 dpi, respectively (Fig. 2C). Interestingly, lymphocytes, including T/NK and B cells, peaked at 16.1% and 16% at 5 dpi, respectively, but experienced a substantial crash to 4.3% and 5.4% at 14 dpi (Fig. 2C). In contrast, myeloid cells such as monocytes, macrophages, dendritic cells, collectively referred to as “Mono/Mac” increased as the fungal burden increased, starting at 8.6% at 0 dpi and rising to 18.7% at 14 dpi. Neutrophils also displayed a dramatic increase, from 6.6% at 0 dpi to 57.6% at 14 dpi (Fig. 2C).

**Figure 2:**
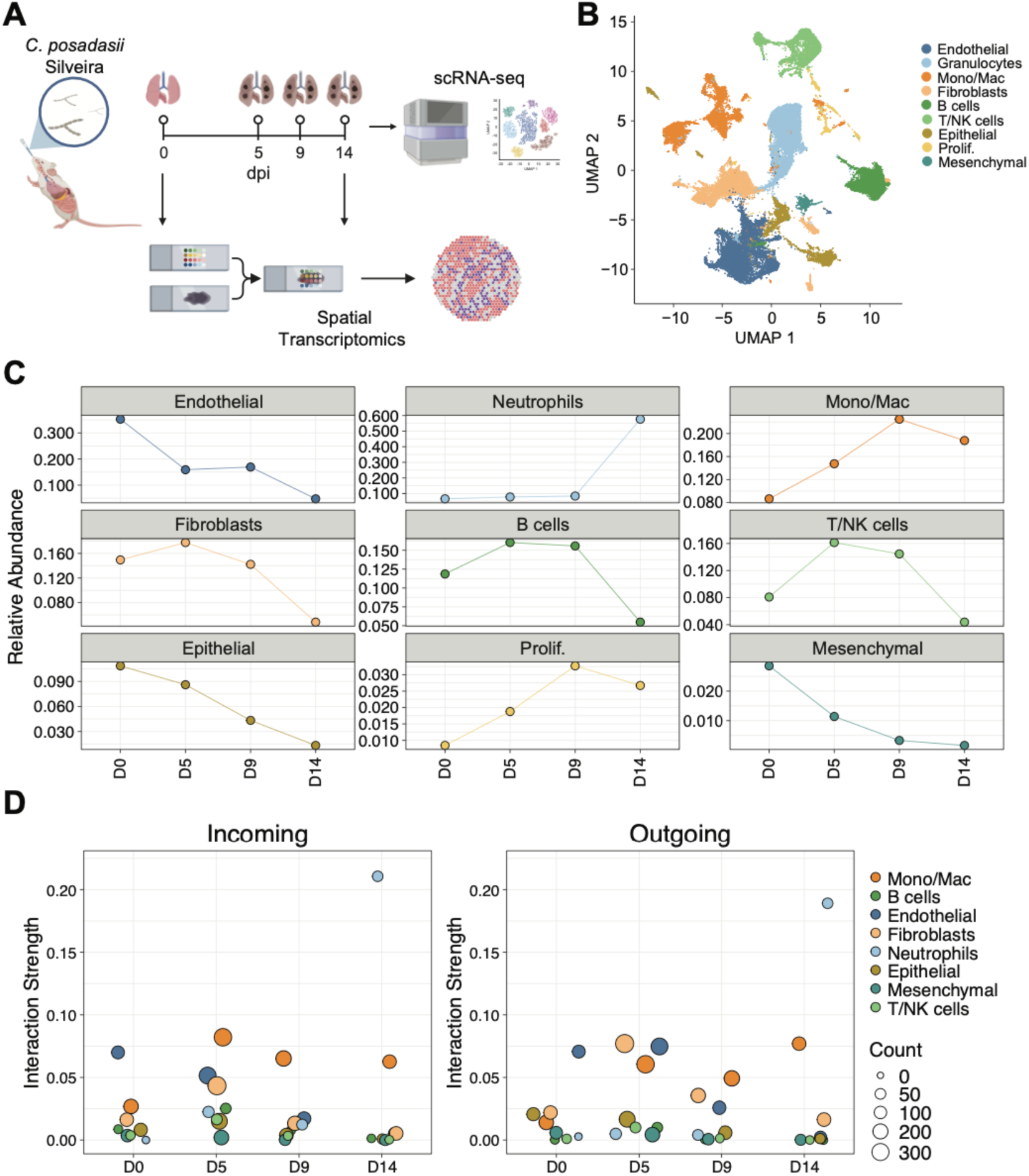
scRNAseq reveals shifts in cell populations and immune infiltration in the lungs following *C. posadasii* infection. (A) Diagram illustrating the experimental design for the transcriptomic experiments. scRNAseq was performed on non-infected (0 dpi) and infected (5, 9, 14 dpi) lungs from mice that were intranasally infected with 384-540 *C. posadasii* Silveira. Each time point represents a pooled sample of n=3. Spatial transcriptomics was performed at 0 dpi and 14 dpi lungs. Created with BioRender.com (B) UMAP plot of the main cell types of infiltrating and resident cells in the lung from non-infected (0 dpi) and infected (5, 9, 14 dpi) samples. Cells are color-coded by the main cell types. (C) Faceted line plot depicting the relative abundance of each cell type in relation to all cells. Each subplot corresponds to a specific cell type and is color-coded accordingly. The X-axis represents the time points, while the Y-axis indicates the relative abundance. (D) Dot plot illustrating the inferred incoming and outgoing interaction signaling strength. The left plot represents the incoming interaction strength, while the right plot represents the outgoing interaction strength. The X-axis denotes the time points, and the Y-axis indicates the interaction strength. The color of each dot corresponds to a specific cell type, and the size of the dot reflects the number of inferred interactions for that cell type at each time point.

Analysis of cellular interactions revealed that prior to infection, endothelial cells displayed the highest levels of both incoming and outgoing signaling strength. Upon infection, shifts in signaling strength were observed, consistent with an increase in myeloid cell populations and the corresponding decrease in all other cell populations (Fig. 2C, D). At 5 dpi, signaling strength to and from Mono/Mac cells increased both in terms of interaction strength and number of interactions, and remained high throughout the course of infection. By 14 dpi, the signaling landscape was dominated by neutrophils, which were highly abundant at this time point (Fig. 2D). Notably, the only non-immune cell type to increase in both incoming and outgoing signaling strength upon infection were fibroblasts (Fig. 2 D). As the infection progressed, we observed notable shifts in specific signaling pathways, including SPP1, COMPLEMENT, TNF, and TGFb pathways, among others (Fig. S1 D). These results highlight the extensive immune infiltration by myeloid cells, which robustly occupy the lung space and dominate the signaling landscape after infection, with prominent pathways involving inflammatory and fibrotic processes that increase as the infection progresses.

### Infection is associated with a loss of homeostasis and a shift towards a profibrotic phenotype

To further understand the role of non-immune cells (endothelial, epithelial, fibroblasts, mesenchymal) during infection, 15,737 cells across all time points from these clusters were extracted and re-clustered. From the four major cell clusters of non-immune cells, we identified 17 distinct subtypes: alveolar epithelial cell 1 (AT1), alveolar epithelial cell 2 (AT2), club, ciliated, basal, goblet, smooth muscle cells (SMC), lipofibroblasts (Lipofib), mesenchymal alveolar niche cell (MANC), myofibroblasts (Myofib), pericytes, mesothelial, arterial endothelial cells (AEC), venous endothelial cells (VEC), lymphatic endothelial cells (LEC), general capillary cells (gCAP), and alveolar capillary cells (aCAP) (Fig. 3 A, B; Fig. S2 B; and Table S1). Despite the overall decrease in non-immune cells following infection, we observed an increase in several non-immune subpopulations at the early stages of infection, including SMC and mesothelial cells from the mesenchymal lineage, and fibroblast lineage cells such as Myofib and MANC cells (Fig. S2C).

**Figure 3:**
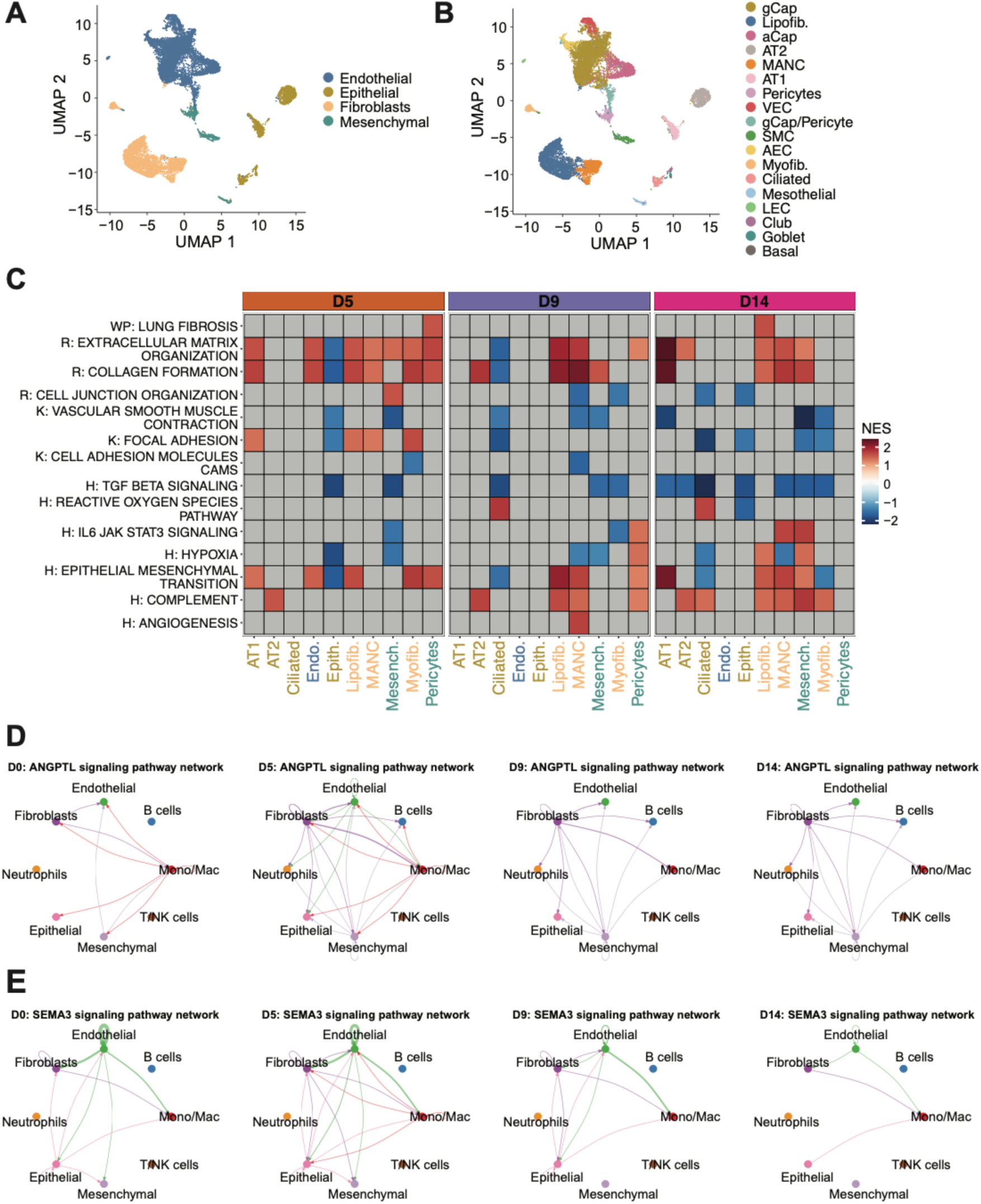
Non-immune cells shift towards a profibrotic phenotype. (A) Integrated UMAP plot of the main non-immune cell types in the lung from uninfected infected samples. Cells are color-coded by identified cell type. (B) Integrated UMAP plot of subclustered non-immune cell types in the lung. Cells are color-coded by identified subtype. (C) Heatmap of pathway activities for non-immune cell types, scored by Gene Set Enrichment Analysis (GSEA) of differentially expressed genes. Pairwise differential expression comparisons were performed between each indicated timepoint and 0 dpi. Genes were ranked by their -log10 p-value, multiplied by the sign of the log2 fold change, and analyzed using the fgsea package. Enrichment is represented by the normalized enrichment score (NES), with blue indicating significant downregulation, red indicating significant upregulation, and gray representing non-significant NES values (FDR ≥ 0.05). Pathways from MSigDB (KEGG, Reactome, Hallmark, and WikiPathways) were curated and abbreviated as K, R, H, and WP, respectively. The X-axis denotes the cell type, and the Y-axis denotes the pathway. (D) Circle plots of ANGPTL signaling pathway network, with separate plots for each time point. Colors correspond to the main cell types. In each plot, edge colors represent the sources as the sender, and edge weights correspond to the interaction strength, with thicker lines indicating a stronger signal. (E) Circle plots of SEMA3 signaling pathway network, with separate plots for each time point. Colors correspond to the main cell types. In each plot, edge colors represent the sources as the sender, and edge weights correspond to the interaction strength, with thicker lines indicating a stronger signal.

Next, we performed a functional/pathway enrichment analysis by identifying differentially expressed genes at each time point compared to uninfected cells. Throughout the course of the infection, fibroblasts and mesenchymal cells exhibited a consistent increase in enrichment for lung fibrosis pathways, including Epithelial-Mesenchymal Transition, Collagen Formation and Extracellular Matrix Organization (Fig. 3C). After 5 dpi, fibroblasts and mesenchymal cells also showed enrichment for the Complement pathway. Supporting these enriched pathways, cell communication analysis identified fibroblasts as the dominant cell type associated with the ANGPTL and IL6 signaling pathways, which have been linked to lung fibrosis and lung leakage (19–21) (Fig. 3 D and Fig. S2 D). Notably, the primary receivers of the of the ANGPTL and IL6 fibroblast signals were the Mono/Mac cells (Fig. 3 E and Fig. S2 D). Interestingly, we also observed a strong inferred signaling of the Oncostatin M (OSM) pathway, a cytokine from the IL6 family (22), from Mono/Macs to fibroblasts (Fig. S2E). Epithelial cells exhibited varied enrichment patterns. AT1 epithelial cells showed enrichment for the same fibrosis pathways observed in fibroblasts at 5 dpi. In contrast, AT2 epithelial cells consistently demonstrated enrichment for the Complement pathway. Lastly, endothelial cells exhibited a peak enrichment at 5 dpi for fibrosis-related pathways, including Epithelial-Mesenchymal Transition, Collagen Formation, and Extracellular Matrix Organization. No further enrichment was observed throughout the remainder of the infection. These findings are summarized in Fig. 3 C. The SEMA3 signaling pathway, associated with lung homeostasis (23, 24) is driven by endothelial cells and decreased as the infection progressed (Fig. 3 D). These results highlight a lung in a state of dysfunction, with homeostatic signaling decreasing over time and a shift towards a profibrotic phenotype.

### Lymphocytes fail to mount a robust anti-fungal response

Lymphocytes, in particular T cells, are known to play a crucial role in the defense against *Coccidioides spp.* in both human and mice (11–16). While lymphocytes initially increased at 5 dpi in our data set, they subsequently declined to a proportionally lower population size than before the infection by 14 dpi (Fig. 1 C). To further characterize the response of lymphocyte populations during infection, we extracted and re-clustered all lymphocyte clusters (T/NK cells, B cells) using an unsupervised clustering approach. 8659 lymphocytes were analyzed in total across all time points and were further subdivided into 9 subtypes: B cells, CD8 T, CD4 T, Natural Killer (NK), Innate lymphoid cells (ILCs), Gamma delta T (gdT), regulatory T cells (Tregs), Interferon-Stimulated Gene-expressing (ISG) B cells, Natural killer T (NKT) (Fig. 4 A, C; and Table S1). In assessing the relative abundance of these subpopulations, we observed that B cells, the largest lymphocyte subpopulation, peaked at 9 dpi at 11.4% and subsequently declined to 4% by 14 dpi. The next largest subpopulations were CD8 T, CD4 T, and NK cells, all peaking at 5 dpi at 4.7%, 4.1%, and 2.9%, respectively. These cells decreased substantially to 1%, 0.97%, and 0.27% by 14 dpi. The remaining subpopulations were relatively small and ended with fewer cells than they started with by 14 dpi. The exception was Tregs, which started at 0.16% and increased to 0.48% by 14 dpi. These results are summarized in Fig. 4 B.

**Figure 4:**
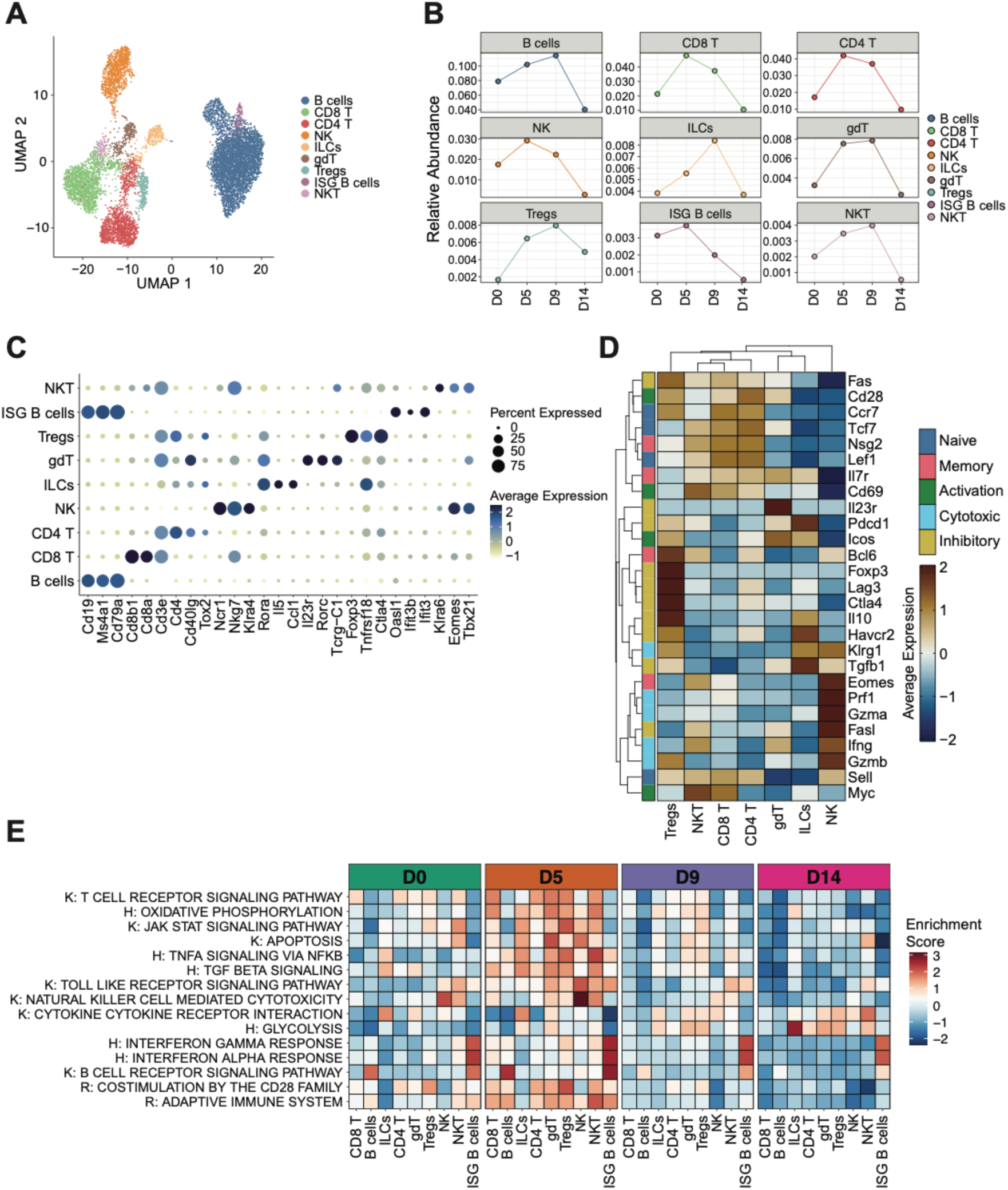
Lymphocytes fail to mount robust anti-fungal response. (A) Integrated UMAP plot of lymphocyte cell types in the lung from uninfected (0 dpi) and infected (5, 9, 14 dpi) samples. Cells are color-coded by identified cell types. (B) Faceted line plot depicting the relative abundance of each cell type in relation to all cells. Each subplot corresponds to a specific cell type and is color-coded accordingly. The X-axis represents the timepoints, while the Y-axis indicates the relative abundance. (C) Dot plot illustrating the average expression levels of select genes. Expression levels are color-coded from low (muted yellow) to high (deep indigo). The size of each circle represents the percentage of cells expressing the gene. (D) Heatmap illustrating the scaled average expression levels of selected T cell genes. Expression levels are color-coded from low (dark blue) to high (dark brown). Column names represent the lymphocyte cell types excluding B cells, with colors on top of each column indicating their respective groups. Row names correspond to the selected genes. Both genes and subtypes are hierarchically clustered based on expression similarity. (E) Heatmap of pathway activities in lymphocytes scored per cell by AUCell. Enrichment is represented by a scaled average AUCell score per cell type, with blue indicating lower enrichment and red indicating higher enrichment. Pathways from MSigDB (KEGG, Reactome, Hallmark, and WikiPathways) were curated and abbreviated as K, R, H, and WP, respectively. The X-axis denotes the cell type, and the Y-axis denotes the pathway.

Both CD4 and CD8 T cells peaked in terms of cell proportions and enrichment scores at 5 dpi, indicating an initial expansion and potential transition to an active state in response to infection. This is highlighted by the pathways Oxidative Phosphorylation (OXPHOS), T Cell Receptor Signaling Pathway, and Costimulation by the CD28 Family (Fig. 4 E). We observed high expression of classic marker genes for naïve and activated T cells in the two populations (Fig. 4 D). This observation is supported by pathway enrichment analyses at 0 and 5 dpi, which suggest that both T cell populations are mixed, containing both naïve and activated cells early during the infection. Notably, we observed a considerable drop off in the activity of both CD4 and CD8 T cells at 9 and 14 dpi (Fig. 4 E). Tregs showed early enrichment for OXPHOS at 5 dpi, indicating early suppressive capacity and stability (Fig. 4 E) (25, 26). Additionally, classical inhibitory genes (*Il10, Fas, Ctla4, Gzmb*) were highly expressed in Tregs (Fig. 4 D).

To further characterize the CD4 and CD8 T cell populations, we isolated each cell type and subclustered independently. Our analysis identified two subpopulations of CD4 T cells: Naïve and Th17 (*S100a11, S100a4*) (27), with *Il17* expression not detected in the Th17 cells (Fig. S3 C). Using pseudotime trajectory analysis, we identified a single lineage where naïve cells converged on the Th17 subpopulation (Fig. S3 C). As the CD4 trajectory progressed we see a shift from the naïve-like expression profile towards an inhibitory/regulatory expression profile (Fig. S3 C). For CD8 T cells, we identified three subtype populations: naïve, effector, and exhausted. Trajectory analysis revealed a single lineage where naïve and effector cells converged on the exhausted subpopulation, with overlapping patterns of gene expression apparent between effector and exhausted cells (Fig S3 D).

NK cell numbers also peaked at 5 dpi, with Natural Killer Cell Mediated Cytotoxicity as the top enriched pathway, followed by a subsequent decline (Fig. 4B, E). NK cells exhibited the highest expression of canonical cytotoxic genes (*Klrg1, Gzma, Gzmb, Prf1*). The smaller populations of ILCs, gdT, and NKT cells showed enrichment for signaling pathways such as Jak-STAT Signaling Pathway, TNFA Signaling Pathway, and TGF Beta Signaling, along with the expression of some inhibitory genes. These findings are illustrated in Fig. 4 D and E. Lastly, enrichment for the B Cell Receptor Signaling Pathway was observed within B cells, which peaked early at 5 dpi and subsequently declined over time, with no other pathways being enriched during the infection (Fig. 4 E). Taken together, these results highlight an ineffective adaptive response during *C. posadasii* infection, characterized by an early expansion followed by a sharp decline in most lymphocyte populations as the infection worsens.

### Diverse myeloid responses during infection reveal a large expanding profibrotic macrophage population

Next, we investigated the response of myeloid cells, which showed a significant expansion at 5 dpi (Fig. 1C). The “Mono/Mac” cell cluster, comprised of monocytes, macrophages, and dendritic cells, totaling 5291 cells across all time points, was extracted and reclustered, revealing the following 11 subtypes: Alveolar Macrophage (AM), Spp1^+^ Macrophage (Spp1^+^ Mac), Classical Monocyte (C Mono), Non-Classical Monocyte (NC Mono), Dendritic cell (DCs), MHCII Interstitial Macrophage (MHCII IM), Prolif, Lyve1^+^ Interstitial Macrophage (Lyve1^+^ IM), Cd5l^+^ Macrophage (Cd5l^+^ Mac), migratory Dendritic cell (migDC), ISG-expressing Monocyte (ISG Mono) (Fig. 5 A; Fig. S4 B; and Table S1). In assessing the relative abundance of the subpopulations, we observed that the largest population, AM, initially increased but eventually returned to proportions similar to those at 0 dpi (Fig. S4 C). The second largest population, the Spp1^+^ Mac population, showed a significant increase, rising from no cells observed at 0 dpi to 7.5% of the total lung at 14 dpi (Fig. S4 C). Both monocyte populations, C Mono and NC Mono, decreased from their initial abundances of 1% and 1.3% to 0.9% and 0.8%, respectively (Fig. S4 C). The DC populations, although relatively small, initially increased in abundance but by 14 dpi, both had smaller populations than before the infection (Fig. S4 C). The interstitial macrophage populations, MHCII IM and Lyve1^+^ IM, showed increases in their relative abundances, rising from 0.26% and 0.1% to 0.81% and 0.91%, by the end of the infection (Fig. S4C). The relatively small and distinct Cd5l^+^ Mac population exhibited a notable increase, reaching 1.8% at 14 dpi (Fig. S4 C).

**Figure 5:**
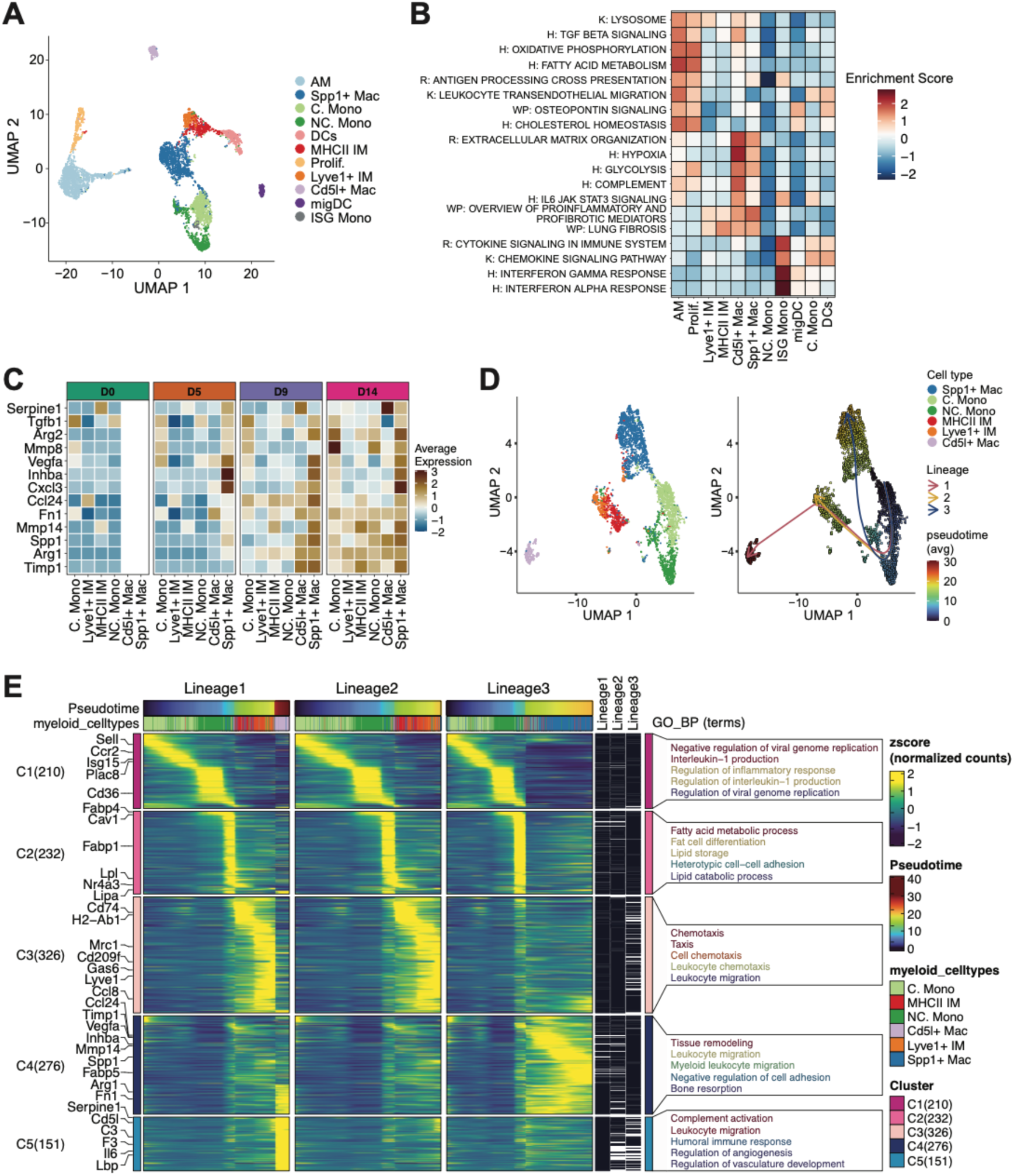
Infection-Driven Expansion of Profibrotic Macrophages. (A) Integrated UMAP plot of Mono/Mac cell types in the lung from uninfected (0 dpi) and infected (5, 9, 14 dpi) samples. Cells are color-coded by identified cell types. (B) Heatmap of Mono/Mac cell type pathway activities scored per cell by AUCell. Enrichment is represented by a scaled average AUCell score per cell type, with blue indicating lower enrichment and red indicating higher enrichment. Pathways from MSigDB (KEGG, Reactome, Hallmark, and WikiPathways) were curated and abbreviated as K, R, H, and WP, respectively. The X-axis denotes the cell type, and the Y-axis denotes the pathway. (C) Heatmap illustrating the scaled average expression levels of selected fibrosis-related genes. Expression levels are color-coded from low (dark blue) to high (dark brown). The column names represent a subset of Mono/Mac cell types, excluding dendritic cells, alveolar macrophages, and proliferating cells. The colors on top of each column indicate their respective groups. The row names correspond to the selected genes. Both genes and cell subtypes are hierarchically clustered based on expression similarity. (D) UMAP plot displaying pseudotime trajectories. Cells are color-coded based on their pseudotime values, ranging from blue (early) to red (late). The pseudotime trajectory was inferred in an unsupervised manner using Slingshot. Each arrow represents a distinct identified trajectory (lineage). (E) Heatmap illustrating the scaled fitted gene expression values along pseudotime. Expression levels are color-coded from low (purple) to high (yellow). Column-wise breaks in the heatmap separate each lineage (pseudotime trajectory) into distinct panels. The top row for each lineage represents pseudotime, ordered from left to right and color-coded based on pseudotime values, ranging from blue (early) to red (late). The next row contains cell type colors, where each bar represents a cell belonging to a specific cell type along the pseudotime trajectory. Rows represent genes, with selected genes highlighted on the left-hand side. Genes were clustered based on fitted expression values, with each cluster represented by row-wise breaks in the heatmap. Each cluster is labeled on the left-hand side by “C#” followed by the number of genes in that cluster in parentheses. Different gene clusters can be distinguished by the clusters on the left-hand side of the heatmap, represented by a color block. Genes from each cluster were then analyzed for gene ontology (GO) enrichment, with the top 5 enriched terms for each cluster shown on the right side of the heatmap.

We observed enrichment for various metabolic, immune, and tissue remodeling pathways, such as Lysosome, TGF-beta signaling, Oxidative Phosphorylation, and Osteopontin signaling in the AM population (Fig. 5 B). Spp1^+^ Mac and Cd5l^+^ Mac exhibited enrichment for inflammatory, hypoxic, and fibrotic pathways, including Hypoxia, Complement, Overview of Proinflammatory and Profibrotic Mediators, and Lung Fibrosis (Fig. 5 B). A key difference between Spp1^+^ and Cd5l^+^ Mac was the greater enrichment for Hypoxia, Complement, and IL6 JAK-STAT3 Signaling in Cd5l+ Mac. Both interstitial macrophages showed slight enrichment for Overview of Proinflammatory and Profibrotic Mediators and Lung Fibrosis (Fig. 5 B). NC Mono did not show enrichment for any pathways. In contrast, C Mono and DCs were enriched for signaling pathways such as Chemokine Signaling Pathway and Cytokine Signaling in the Immune System (Fig. 5 B).

Spp1^+^ macrophages were further characterized due to their dramatic expansion and recent association with various pathological conditions such as severe infection, fibrosis, and poor prognosis in cancer (28–33). Consistent with their increased numbers at late-stage infection, large accumulation of cells expressing Osteopontin (OPN), the protein produced by the *Spp1* gene, was observed within the lungs at 14 dpi (Fig S4 F). We removed the Prolif, DC, and AM populations and reprocessed the remaining cells together (Fig. 4 D, S4 D). The Spp1^+^ Mac cluster exhibited increased expression of fibrosis-associated genes, consistent with its enrichment in the lung fibrosis pathway (Fig. 5 B and C). Pseudotime analysis revealed three distinct lineages emerging from C Mono. Lineage 1 included C Mono, NC Mono, MHCII IM, Lyve1^+^ IM, and Cd5l^+^ Mac. Lineage 2 consisted of C Mono, NC Mono, MHCII IM, and Lyve1^+^ IM. Finally, lineage 3 compriseed C Mono, NC Mono, and Spp1+ Mac (Fig. 5 D and Fig. S4 E). For each lineage, we observed a transition from monocytes to monocyte-derived macrophages. Lineage 2 culminates in the generation of MHCII IM and Lyve1^+^ IM. Notably, the Lyve1^+^ IM population acts as a progenitor for the Cd5l^+^ Mac population, which is the endpoint of lineage 1. Lineage 3 is the most distinct, as it shows a direct transition from monocytes to the Spp1^+^ Mac population. Analysis of the expression patterns along these lineages revealed five distinct clusters of gene expression. Clusters 1 and 2 (C1, C2) largely consisted of genes highly expressed in C Mono and NC Mono populations, showing an enrichment for interleukin-1 production, antiviral response, and lipid production (Fig. 5E). C3 exhibited an enrichment for chemotaxis and leukocyte migration, with high expression of genes primarily found in the MHCII IM and Lyve1^+^ IM populations (Fig. 5E). C4 displayed high expression of genes found in the Spp1^+^ Mac population, with an enrichment for migration and tissue remodeling (Fig. 5E). Lastly, C5 showed high expression of genes found in the Cd5l^+^ Mac population, with an enrichment for complement activation and migration (Fig. 5E). Collectively, these findings highlight the diverse myeloid response with complex cellular differentiation pathways during *C. posadasii* infection, including the identification of a profibrotic macrophage population extensively associated with poor outcomes.

### Extensive infiltration of diverse neutrophil populations with varied inflammatory properties during late-stage infection

All neutrophils from the initial cluster (3542 cells) were extracted and reclustered into 5 neutrophil subtypes: Circulating Neutrophils (Circulating Neu), Early Neutrophils A (Early Neu A), Early Neutrophils B (Early Neu B), ISG-expressing Neutrophils (ISG Neu), PD-L1^+^ Neutrophils (PD-L1^+^ Neu) (Fig. 6 A; Fig. S5 C; and Table S1). Circulating Neu were present at 0 dpi (Fig. S5 A, D); their numbers increased at 5 dpi, peaking at 3.9%, before decreasing towards the latter stages of the infection (Fig. S5 D). ISG Neu, although a small population, showed a steady increase throughout the infection, rising from 0.07% to 0.97% (Fig. S5 D). Both Early Neu A and B, as well as the PD-L1^+^ Neu populations exhibited a dramatic increase at 14 dpi, with Early Neu A and B increasing to 2.1% and 11.9%, respectively, and PD-L1^+^ Neu to 9.3% (Fig. S5 D). The increase in neutrophils, as well as progressive upregulation of PD-L1 on the cell surface, was confirmed by flow cytometric analysis (Fig. 6 B and Fig S5 B). The PD-L1 Neu cluster exhibited enrichment TNF, Complement, and Inflammatory response as well as Hypoxia and Apoptosis pathways (Fig. 6 C). This is consistent with the cell-cell communication analysis, which indicates that neutrophils have high signaling activity in TNF and Complement pathways at 14 dpi (Fig. 6C and Fig. S5 F and G). Circulating Neu demonstrated strong enrichment in migratory pathways, such as Cell Surface Interactions at the Vascular Wall and Leukocyte Transendothelial Migration (Fig. 6 C). The Early Neu A and B clusters were surprisingly distinct, with Early Neu A showing high enrichment for Neutrophil Degranulation, Reactive Oxygen Species, and Neutrophil Extracellular Trap Formation, whereas Early Neu B exhibited only slight enrichment in the E2F Targets pathway (Fig.6 C). Paradoxically, many genes expressed in both Early Neu A and B are those typically expressed during early neutrophil development within the bone marrow, hence the nomenclature ‘Early Neu’ (34, 35).

**Figure 6:**
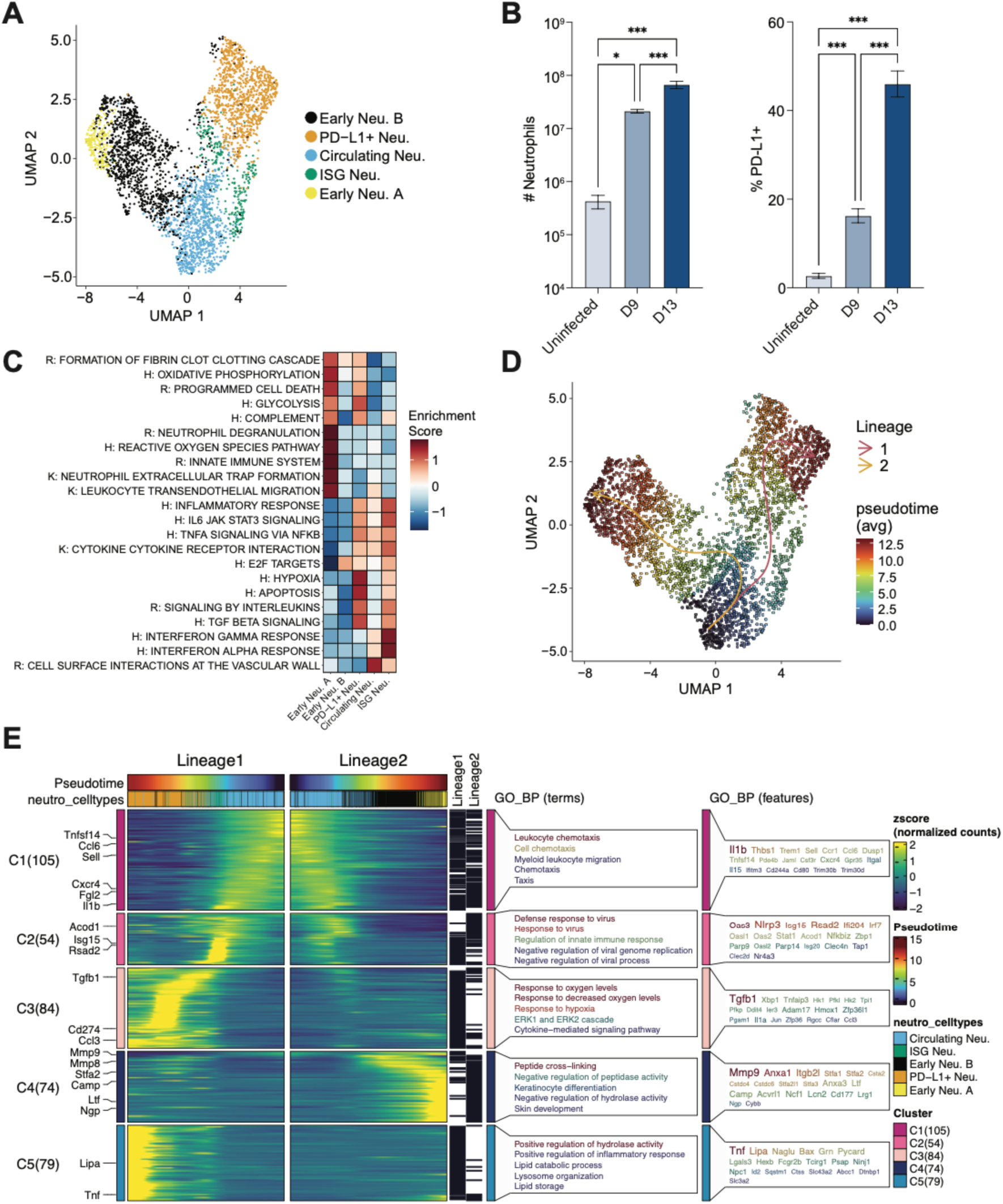
Diverse Neutrophil Response at Late-Stage Infection. (A) An integrated UMAP plot of subclustered neutrophil cell types in the lung from uninfected (0 dpi) and infected (5, 9, 14 dpi) samples. Cells are color-coded by identified subpopulation. (B) Quantification of total and PD-L1^+^ neutrophils in the lungs. Bar graphs represent the mean (n=6) +/- standard deviation. C) Heatmap of neutrophil pathway activities scored per cell by AUCell. Enrichment is represented by a scaled average AUCell score per cell type, with blue indicating lower enrichment and red indicating higher enrichment. Pathways from MSigDB (KEGG, Reactome, Hallmark, and WikiPathways) were curated and abbreviated as K, R, H, and WP, respectively. The X-axis denotes the cell type, and the Y-axis denotes the pathway. (D) UMAP plot displaying pseudotime trajectories. Cells are color-coded based on their pseudotime values, ranging from blue (early) to red (late). The pseudotime trajectory was inferred in an unsupervised manner using Slingshot. Each arrow represents a distinct identified trajectory (lineage). (E) Heatmap illustrating the scaled fitted gene expression values along pseudotime. Expression levels are color-coded from low (purple) to high (yellow). Column-wise breaks in the heatmap separate each lineage (pseudotime trajectory) into distinct panels. The top row for each lineage represents pseudotime: lineage 1 is ordered from right to left (red to blue), and lineage 2 from left to right (blue to red). The next row contains cell type colors, where each bar represents a cell belonging to a specific cell type along the pseudotime trajectory. Rows represent genes, with selected genes highlighted on the left-hand side. Genes were clustered based on fitted expression values, with each cluster represented by row-wise breaks in the heatmap. Each cluster is labeled on the left-hand side by “C#” followed by the number of genes in that cluster in parentheses. Different gene clusters can be distinguished by the clusters on the left-hand side of the heatmap, represented by a color block. Genes from each cluster were then analyzed for gene ontology (GO) enrichment, with the top 5 enriched terms and genes for each cluster shown on the right side of the heatmap. *p<0.05, ***p<0.001.

Trajectory analysis revealed two distinct lineages originating from Circulating Neu. The first lineage progressed from Circulating Neu to ISG Neu and culminated in PD-L1^+^ Neu (Fig. 6 D). The second lineage also began with Circulating Neu but diverged towards Early Neu B and ultimately transitioned to Early Neu A (Fig. 6 D). Expression analysis along these lineages revealed 5 expression clusters. C1 consisted of genes highly expressed in Circulating Neu, with an enrichment for chemotaxis and migration (Fig. 6E). C2 showed an enrichment for response to viruses, with this cluster having high expression of genes primarily found in the ISG Neu cluster (Fig. 6E). C3 and C5 exhibited high expression of genes found in the PD-L1^+^ Neu population, with an enrichment for hypoxia and inflammatory responses (Fig. 6E). Lastly, C4 showed high expression of genes found in both Early Neu A and B, with an enrichment for peptide cross-linking and negative regulation of peptidase and hydrolase activities, suggesting a potential regulation of NETosis (36) (Fig. 6E). Taken together, these data highlight the dramatic and extensive infiltration of diverse neutrophil populations, which exhibit a wide range of inflammatory characteristics during the later stages of infection.

### Significant reorganization of lung cellularity and molecular pathways during infection

Spatial transcriptomics was utilized to visualize lung cellularity and spatial organization of cell populations comparing uninfected and end stage infection samples (14 dpi; Fig. 2A and Fig. 7A). At 0 dpi, the lung was primarily composed of endothelial, epithelial, and fibroblast populations with an underrepresentation of immune cells (Fig. 7B). However, at 14 dpi, there was a notable decrease in endothelial, epithelial and mesenchymal cells (Fig. 7C). Strikingly, at 14 dpi, we observed large regions of dense cellular infiltrates, consistent with necrotizing granuloma formation (Fig. 7A) (37) along with massive neutrophil infiltration in the same region, supporting the scRNAseq data (Fig. 7C). The Mono/Mac cell population also increased after infection, further corroborating previous findings. Interestingly, we observed colocalization of Mono/Mac and fibroblasts, which were shown in our scRNAseq data to communicate strongly during infection (Fig. 3E and Fig. S2D). These data are summarized in Fig. 7B and C. Expression of neutrophil marker genes *S100a8* and *S100a9* at 0 dpi was spotty and lacked localization, coupled with low expression (Fig. 7G). In contrast, at 14 dpi, both *S100a8* and *S100a9* showed high spatial localization and were highly expressed in the same regions where deconvolution highlighted the neutrophils (Fig. 7G). Interestingly, regions with little to no gene expression were observed at the center of neutrophil dense regions, again consistent with areas of abundant necrosis (Fig. 7G and H). Immunohistochemical analysis further indicated that neutrophils were localized in a distinctive spherical pattern, consistent with accumulation around fungal spherules (Fig. 7D-E, S1B). Other relevant genes, such as *Cd274* (PD-L1) and *Spp1*, did not show spatial localization and had low expression at 0 dpi (Fig. 7G). After infection, *Cd274* and *Spp1* showed increased expression and localization, with *Spp1* co-localizing to regions enriched for fibroblasts (Fig. 7G). Consistent with our scRNAseq data, we observed enrichment of pathways such as TNFA Signaling via NFKB, Complement, and Hypoxia in neutrophil-rich regions in the infected lung (Fig. 7H and Fig. S5I). Additionally, fibrosis-related pathways, including Epithelial Mesenchymal Transition, Lung Fibrosis, and Collagen Formation, were enriched in regions consisting of both Mono/Mac and fibroblast populations (Fig. 7H and Fig. S5H). These findings demonstrate the significant reorganization of lung cellularity and molecular pathways during infection, with large areas of inflammatory neutrophil accumulation around necrotic tissue and co-localized areas of fibroblasts and Mono/Mac cells expressing fibrosis related genes, offering insights into the underlying mechanisms of lung pathology.

**Figure 7:**
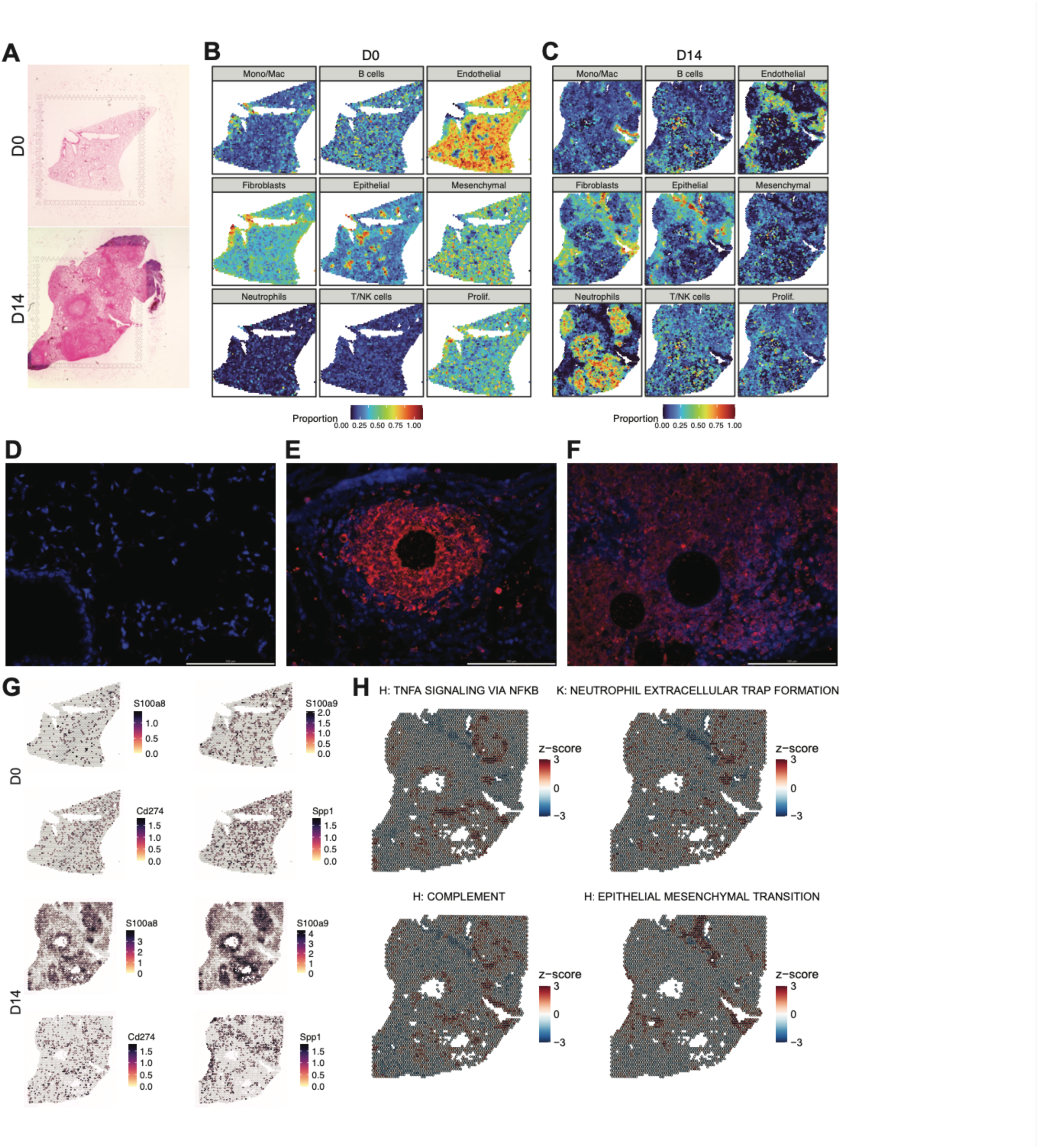
Infection-Driven Reorganization of Lung Cellular and Molecular Dynamics. (A) H&E stained tissue sections used for spatial sequencing at 0 dpi (top) and 14 dpi (bottom). Spatial plots of deconvolution of a 0 dpi (B) or 14 dpi (C) lung using scRNAseq. Each subpanel represents a major cell type from the scRNAseq dataset, with the color of the spots indicating the estimated cell type proportion, ranging from blue (low) to red (high). Merged IHC images of the lungs at 0 dpi (D) or 14 dpi (E and F). DAPI staining is represented in blue, S100A8 is represented in red. The white scale bar in the bottom right-hand corner represents 100 µm. Image taken at 40x magnification. (G) Spatial plot of selected gene expression at 0 dpi (top) and 14 dpi (bottom). Expression levels are represented as SCT normalized counts, with blue indicating low expression and red indicating high expression. (H) Spatial plots of gene set co-regulation analysis (GESECA) on 14 dpi lung for selected pathways. A high positive z-score (red) indicates that the gene set is significantly enriched, while a low or negative z-score (blue) suggests that the gene set is not significantly enriched or has low expression. Pathways from MSigDB (KEGG, Reactome, Hallmark, and WikiPathways) were curated and abbreviated as K, R, H, and WP, respectively.

## Discussion

Despite significant advances, our understanding of the basic mechanisms of fungal pathogenesis within the lungs remains limited, hindering the development of effective vaccines and therapies. To bridge this knowledge gap, we performed a comprehensive spatio-temporal transcriptomic characterization of disease progression in the lungs over time using a mouse model of lethal *C. posadasii* infection. We temporally dissected the heterogeneity of the response of stromal as well as resident and infiltrating immune cells in the lungs over a time course of infection using scRNAseq. We then employed spatial transcriptomic analysis during severe disease to investigate the spatial organization of distinct populations within the lungs.

Our results highlight a progressively dysfunctional immune response marked by limited lymphocyte activation, the emergence of monocyte-derived macrophages enriched in fibrotic signaling pathways, and massive accumulation of diverse pro-inflammatory neutrophil populations. Similarly, the response of non-immune cells in the lung suggests initial interactions with the fungus do not lead to sustained protective responses. Most pathways enriched in non-immune cells are related to fibrosis, including within non-fibroblast cell types. Harding et. al (2024, submitted for publication) report enrichment for pro-inflammatory responses, such as hypoxia and several chemotaxis pathways, in human airway epithelial cell cultures in response to *C. posadasii* infection. The absence of such pro-inflammatory responses in non-immune cells in our study may be contributing to poor infection outcomes, although future studies are needed to confirm this hypothesis.

CD4 T cells play a well-documented role in combating *Coccidioides* in humans and mice (11–16, 38), with protective responses defined mostly in the context of vaccination studies. In this lethal model of *Coccidioides* infection, we observe an initial expansion of lymphocytes followed by a rapid decline, particularly in T and NK cells. Our data suggest impairment in the metabolic reprogramming of T cells. OXPHOS is the dominant metabolic process in naïve/quiescent T cells, which is followed by a well-documented metabolic shift towards glycolysis as T effector cells develop (39, 40). We observe that both CD4 and CD8 T cells show peak enrichment for OXPHOS at 5 dpi followed by a subsequent decrease, although with no corresponding increase in glycolysis at later timepoints. The lack of significant glycolysis enrichment in CD8 T cells and the minimal increase in CD4 T cells at 14 dpi may indicate a compromised effector function, which could contribute to the observed decline in T cell populations. Expansion of Tregs may also be contributing to the decline in both T cell populations. In pediatric patients, elevated Tregs have been associated with chronic *Coccidioides* infections (18). Interestingly, Tregs do exhibit a shift from OXPHOS to glycolysis after 5 dpi along with a steady increase in their relative proportion, indicating activation and a possible role of this subset in inhibiting naïve and activated T cells, reducing overall T cell population sizes and worsening infection outcomes.

A key feature of our data is the striking emergence of Spp1^+^ macrophages. These macrophages, characterized by their expression of *Spp1* and fibrosis related genes, have been linked to severe/poor outcomes in varied disease states, such as severe COVID-19, lung fibrosis, and cancer (lung, colorectal) (29–33). In our study, we observed progressive infiltration of monocytes which ultimately transition to one of three macrophage subsets, the most abundant of which are the Spp1^+^ macrophages. Spp1^+^ macrophages display a distinct fibrotic expression profile, characterized by the upregulation of several key genes, including *Tgfb1*, *Fn1*, *Inhba*, and various matrix metalloproteinases (MMPs), which are known to play crucial roles in tissue remodeling and fibrosis (41–44). Notably, we detected various fibrosis-related signaling pathways with strong inferred interactions between Mono/Mac, fibroblasts, and neutrophils over the course of the infection, including reciprocal signaling of the IL-6 and OSM pathways between Mono/Macs and fibroblasts, which has been implicated in fibrosis and inflammation (22, 45, 46). Interestingly, in our spatial data we observe overlapping localization of Mono/Macs and fibroblasts with enrichment for fibrotic signaling pathways, suggesting active sites of crosstalk between these populations *in vivo* resulting in a fibrotic phenotype.

Intriguingly, in our companion study (Miranda et al 2024, submitted for publication), Spp1^+^ macrophages emerge 24 hours post infection with the attenuated *C. posadasii* Δ*cts2*/Δ*ard1*/Δ*cts3* in a manner that is dependent upon contact with the fungal spore. In this present study, Spp1^+^ macrophages are detectable on 5 dpi but do not robustly emerge until 9 dpi. A key difference between these two studies, in addition to strain selection, is the size of the initial inoculum (500 versus 1×10^6^ arthroconidia). The early emergence of Spp1^+^ macrophages in Miranda et. al may be facilitated by the large inoculum size, whereas in this present study, fungal spread via replication of spherules and endospores may be required to reach sufficiently high fungal burden to induce large numbers of Spp1^+^ macrophages. This further suggests that contact with spherules and/or endospores, in addition to arthroconidia, may be capable of inducing this macrophage subset. Further work is required to clarify which surface molecules on both fungus and host facilitate this differentiation, and furthermore, if this effect is also observed upon contact with human macrophages. Taken together, our two studies highlight a previously undescribed role for Spp1^+^ macrophages in fungal infection, offering new insights into the pathogenesis of *Coccidioides*.

By late-stage infection, neutrophils completely dominate the cellular landscape, representing close to 60% of the total cells in the lung. At early time points, Circulating Neu, which represent standard neutrophils during homeostatic conditions, comprise the most abundant neutrophil population. This population undergoes an expansion in the lungs at 5 dpi, followed by a contraction at 9 dpi and a sharp decrease at 14 dpi in favor of a massive expansion in other neutrophil subsets. Fungal burden in the lungs also exhibits a sharp increase at 14 dpi, suggesting an inability to control fungal replication may help drive the massive neutrophil expansion at this time point. Intriguingly, both Spp1^+^ and CD5L macrophages have previously been implicated in the recruitment and differentiation of neutrophils, specifically PD-L1^+^ neutrophils in the case of Spp1^+^ macrophages (33, 47). Notably, in our dataset, we observed high expression of *Cxcl2* in various macrophage subpopulations, along with strong inferred signaling of the CXCL signaling pathway from Mono/Macs to neutrophils (Fig. S4 H), which has previously been implicated in the recruitment of neutrophils during fibrosis (48). Additional interactions between Mono/Macs and neutrophils were observed via strong inferred signaling from the SPP1 signaling pathway as well as significant crosstalk through the TNF and COMPLEMENT signaling pathways.

Later neutrophil subsets (Early Neu A, B and PD-L1^+^ Neu) have markedly more inflammatory profiles than Circulating Neu. Early Neu A and B were distinguished by their expression of markers for immature and mature neutrophils from the bone marrow (34, 35) and display high granule expression and enrichment for NETosis pathway. The Early Neu subpopulations also exhibit increased expression of *Cd177* and Stefin A family genes, which have all be linked to severe COVID-19 and influenza (36, 49, 50). Similarly, PD-L1^+^ neutrophils have also been observed during severe COVID-19 (33, 51), underscoring a common theme of inflammatory neutrophil association with severe respiratory infections. Similar to previous studies (33), we also observe a strong enrichment for pro-inflammatory signaling and enrichment for the hypoxia pathway in PD-L1^+^ Neu. Our data underscore the complex interplay between neutrophil subpopulations and macrophage signaling during *Coccidioides* infection, highlighting potential parallels with other severe infections and suggesting novel mechanisms of neutrophil regulation that could inform future therapeutic strategies.

Our data suggest a scenario in the lung where uncontrolled fungal replication, potentially due to an ineffective T cell response, induces populations of macrophages with a pro-fibrotic phenotype. Macrophages then signal robustly to neutrophils, inducing massive neutrophil recruitment into the lungs that ultimately develop a pro-inflammatory phenotype, worsening disease outcomes. Alternatively, the emergence of proinflammatory and fibrotic myeloid populations may interfere with the development of appropriate adaptive responses. While speculative, these conclusions offer potential future therapeutic avenues that warrant further investigation. Interventions that target the metabolic profile of T cells, fibrosis inducing cytokines such as IL-6 or OSM, the development of Spp1^+^ macrophages or the secreted OPN protein itself, or the development of inflammatory neutrophils, in particular the PD-L1 molecule which plays a well-known role in suppressing T cells responses in other disease contexts, may all offer the possibility of improved patient outcomes in severe coccidioidomycosis. Future studies that investigate each of these possible therapeutic avenues are paramount.

In conclusion, our comprehensive spatio-temporal transcriptomic analysis of *Coccidioides* infection in the lungs has provided valuable insights into the roles of various immune cell types in disease progression. By utilizing advanced transcriptomic technologies, we have dissected the complex cellular interactions and signaling pathways involved in the immune response during a lethal model of *Coccidioides* infection. These findings not only enhance our understanding of the immune response to *Coccidioides* infection but also suggest potential targets for therapeutic intervention.

## Materials and methods

### Fungal culture

*Coccidioides posadasii* strain Silveira (NR-48944) was obtained through BEI Resources, NIAID, NIH. Arthoconidia cultures were generated as described previously (52). Briefly, liquid 2X glucose 1X yeast Extract (2X GYE) media (VWR, Radnor, PA, USA) was used to grow mycelia cultures in a shaking incubator at 30°C and 150 rpm for 4 days. Mycelia were then streaked onto solid 2X GYE agar and desiccated in an incubator at 30°C for 6 to 8 weeks to allow arthroconidia to develop. Arthroconidia cultures were harvested by scraping growth into 1X PBS (VWR) and filtered through a 70 µM mesh filter. Fungal isolate was then vortexed for 1 min and centrifuged at 12000 x *g* for 20 mins at room temperature. The pellet was resuspended in 30 mL 1X PBS with vortexing and pelleted as before. Final resuspension of the pellet was in 5 mL of 1X PBS. Stocks were stored at 4°C for up to 6 months. Fungal stock and lung titers were assessed by dilution plating on solid 2X GYE agar. All fungal work was performed in Institutional Biosafety Committee approved BSL-3 and ABSL-3 facilities at Lawrence Livermore National Laboratory using appropriate PPE and protective measures.

### Mouse Infections

All animal work was approved by the Lawrence Livermore National Laboratory Institutional Animal Care and Use Committee under protocol #309. All animals were housed in an Association for Assessment and Accreditation of Laboratory Animal Care (AAALAC)-accredited facility. C57BL/6 mice (strain ID 000664) were obtained from Jackson Laboratory (Bar Harbor, ME, USA). 6–10-week-old female mice were used in all experiments.

Mice were infected intranasally with 384-540 (target: 500) arthroconidia in 40 µl (20 µl per nostril) in 1X PBS while under anesthesia (4-5% isoflurane in 100% oxygen). Mice were monitored daily for signs of morbidity and animals were humanely euthanized upon signs of severe disease by CO_2_ asphyxiation. For tissue harvest, animals were anesthetized under isoflurane and the whole animal was perfused with 20 mL sterile PBS containing 50,000 U/L sodium heparin (Sigma) *via* the left ventricle.

### Lung isolation and preparation of single cell suspensions

Following euthanasia and perfusion, lungs were removed and placed in a 100 mm Petri dish containing 1X PBS. A portion of the lung was set aside for fungal burden determination, which was assessed following homogenization in 1X PBS using an Omni Bead Ruptor 4 in 2.8mm ceramic bead tubes at 5m/s for 20s. The rest of the lungs were placed a 100 mm Petri dish containing 5 mL of digestion buffer (DMEM/F12 (Thermo Fisher) + 100 µg/mL DNase I (Roche) and 3 mg/mL collagenase 1 (Worthington)) and minced with scissors into pieces approximately 1 mm in diameter. Minced pieces were then transferred to a 15 mL conical tube and petri dish rinsed with another 5 mL digestion buffer, which was then added to the 15 mL tube. Lung homogenate was incubated for 30 mins in a 37°C shaking incubator at 150 rpm, with cells triturated with a 10 mL pipet at 15 min. At 30 min, supernatant was removed and transferred to a 50 mL conical tube and placed on ice. 10 mL of fresh digestion buffer was then added to remaining lung homogenate and incubated for another 30 min with shaking as above. Following the second 30 min incubation, remaining unhomogenized tissue was pipetted into a 70 µM mesh filter and manually dissociated with the flat end of a pellet pestle. Equal volume DMEM/F12 + 10% fetal bovine serum (FBS, Thermo Fisher) was added to cell suspension to neutralize digestion enzymes and pelleted by centrifugation at 500 x *g* at 4°C for 8 minutes. Red blood cells were removed by incubation in ACK lysis buffer (Thermo Fisher) for 5 min at room temperature. Cells were again pelleted and then resuspended DMEM/F12 + 10% FBS and counted on a Countess II (Thermo Fisher) automated cell counter prior to downstream analysis.

### Histology and Immunohistochemisty

Lungs were collected from infected mice infected at 14 dpi. Following euthanasia and perfusion as described above, the mice were perfused with 20 mL of 10% neutral buffered formalin (NBF). The lungs were then inflated and immersed in 10% NBF at 4°C for 3 days with gentle agitation. Following fixation, the lungs were paraffin-embedded and sectioned into 5 μm thick slices. All sections were dewaxed using xylene and rehydrated with alcohol. Slides were stained following standard hematoxylin and eosin (H&E) staining procedures (Hematoxylin: 3801561, Leica; Bluing Reagent: 6769001, Epredia; Eosin: 3801601, Leica). Samples were cleared using xylene and coverslips were mounted with Permount Mounting Media (SP15-100, Fisher Scientific). Brightfield mages were taken at 10x, 20x, and 40x on a Leica DM4 B microscope. For immunohistochemical staining, heat induced antigen retrieval was performed using Tris-EDTA. To block non-specific binding sites, CAS-block (Thermo Fisher, cat#008120) was applied. Slides were counterstained with Prolong Gold Antifade Mountant with DNA Stain DAPI (Thermo Fisher, cat# P36931) and stained for proteins of interest using the following primary and secondary antibodies: S100A8 (clone: EPR3554, 1:100, Abcam, cat# ab92331), goat anti-rabbit IgG Alexa Fluor 594 (1:500, Thermo Fisher, cat# A11037), OPN (1:100, R&D systems, cat# AF808), and donkey anti-goat Alexa Fluor 594 (1:500, Thermo Fisher, cat# a11058). Immunofluorescent images were taken at 40x on a Leica DM4 B microscope.

### Flow Cytometry

Cells were incubated for 30 min on ice in 100 μl Hank’s balanced salt solution (Thermo Fisher) + 2% FBS with Fc block (1:100 dilution, clone 2.4G2; BD Biosciences) along with the following antibodies, each diluted 1:500: CD45 e450 (clone 30-F11; Invitrogen), CD11b BV510 (M1/70; BioLegend), CD11c PE-Dazzle 594 (clone N418; BioLegend), Ly-6G BV711 (clone 1A8; BD Biosciences), and CD274 (PD-L1) AF488 (clone MIH5; Invitrogen). Cells were then fixed using BD Cytofix/Cytoperm (BD Biosciences) according to manufacturer’s instructions. Flow cytometry was performed using a FACSAria Fusion and data were analyzed using FlowJo v10.10 software (BD Biosciences). Neutrophils were resolved as CD45^+^ CD11b^+^ CD11c^-^ Ly-6G^+^ as described previously (53), with sequential gating scheme depicted in Fig S5 B, and the proportion of PD-L1^+^ neutrophils was quantified. Neutrophil counts were calculated from total lung cell counts and cell proportions as determined by flow cytometry.

### Single-cell Library Preparation and Sequencing

Single-cell RNA sequencing was performed using the 10x Genomics Chromium Single Cell Fixed RNA Profiling (Flex) Kit (cat# 100475), following the manufacturer’s protocol. Following the generation of single-cell suspensions as described above, 3×10^6^ cells per lung sample were fixed using 10x Genomics fixation buffers supplemented with paraformaldehyde to a final working solution of 4% according to manufacturer’s instructions and then stored at -80°C in 10x Genomics long-term storage solution prior to library preparation. Each reaction was inclusive of 3 mice validated for fungal infection and pooled at 500,000 cells per mouse. Probe hybridization was then performed overnight followed by pooling before loading onto the Chromium X Controller (10x Genomics) to generate single-cell gel bead-in-emulsions (GEMs). Reverse transcription, cDNA amplification, and library amplification were performed according to the manufacturer’s recommendation. Quality control measures, including quantification and size distribution analysis, were performed using an Agilent Tape Station. Sequencing libraries were then sequenced on an Illumina NextSeq 2000 platform, producing paired-end reads. Data processing, including alignment, barcode assignment, and gene expression quantification, was performed using the Cell Ranger software pipeline (v 7.2.0) from 10x Genomics (Zheng et al., 2017). Alignments were carried out using the reference genome package refdata-gex-mm10-2020-A for GRCm38 (mm10), provided by 10x Genomics. Additionally, the "Chromium_Mouse_Transcriptome_Probe_Set_v1.0.1_mm10_2020-A" probe set from 10x Genomics was utilized.

### Single-cell RNA sequencing (scRNAseq) data analysis

#### Data Preprocessing

Analysis was initially performed using Seurat (v 4.3.0) (54) and R (v 4.3.2) but transitioned to Seurat (v 5.1.0) (55) and R (v 4.4.0). We created Seurat objects using the ‘CreateSeuratObject’ for each sample with default parameters and then merged into a single object. Quality control (QC) of the datasets involved retaining cells with at least 200 and no greater than or equal to 75000 counts, cells with at least 200 and no greater than or equal to 10000 expressed genes, and cells with a percentage of mitochondrial genes per cell less than 10% (200 >= nCount_RNA <= 75000, 200>= nFeature_RNA <= 10000, and percent_mt < 10). Additionally, we excluded lowly expressed genes by keeping only those genes that were expressed in at least 10 cells. Data was normalized using NormalizeData function using default parameters.

#### Initial Broad Analysis

First, using the ‘FindVariableGenes’ function, we identified 3000 Highly variable genes (HVGs). Next, data was scaled with the effects of cell counts and mitochondrial percentages regressed out using ‘ScaleData’ (vars.to.regress=c("percent_mt", "nCount_RNA"). Subsequently, after performing principal component analysis (PCA) using the ‘RunPCA’ function, the top 50 principal components (PCs) were selected for downstream integration, clustering, and dimension reduction.

Data integration was performed using Harmony (v1.2.0) (56) with the grouping variable set to “orig.ident” which contained all samples. Clustering analysis was performed using ‘FindNeighbors’ and ‘FindClusters’, with the reduction parameter set to "harmony" and resolution of 0.2. Subsequently, a non-linear dimensionality reduction was conducted using Uniform Manifold Approximation and Projection (UMAP), with RunUMAP (reduction = "harmony", umap.method = "uwot").

For cell type annotation, we utilized genes differentially expressed between clusters, known marker genes, and the LungMAP (57) scRNAseq resource to distinguish the different cell populations. Briefly, we used FindAllMarkers (only.pos = true) to identify marker genes of the generated clusters. Marker genes that met our filtering criteria (adjusted p-value < 0.05 and average log2 fold change ≥ 1) were retained. Next, we first transferred the labels (lineage_level1, lineage_level2, celltype_level2) from the mouse LungMAP scRNAseq dataset (58) to our data using two functions: FindTransferAnchors (reference = lungmap, query = ourdata) and TransferData (dims = 1:30). Using the identified marker genes and comparing them with known markers genes and the cell type predictions, we annotated the cells accordingly.

#### Analysis of non-immune cell subsets

Analysis of non-immune cell subsets consisted of extracting all cells labeled as “Endothelial”, “Epithelial”, “Fibroblasts”, and “Mesenchymal” from the broad labels assigned during the initial analysis and performing an in-depth analysis. We followed the same approach used for the initial broad analysis with a few modifications. We identified 2000 HVGs. For dimension reduction, integration, and clustering we used 30 PCs and a cluster resolution of 0.5. Cell annotation was performed as previously stated leveraging a combination of marker genes, differential expression, and LungMAP.

#### Lymphocytes

To further analyze the lymphocyte sub-populations, we extracted all cells labeled as “T/NK cells”, and “B cells”, from the broad labels assigned during the initial analysis. We followed the same approach used for the broad analysis with a few modifications. We identified 2000 HVGs. For dimension reduction, integration, and clustering we used 20 PCs and a cluster resolution of 0.5. As previously described, cell annotation was achieved using a combination of marker genes, differential expression analysis, and data from LungMAP. To identify the various T cell subsets, we performed sub clustering separately on the CD4 and CD8 T cell-labeled populations. We leveraged Slingshot (v 2.12.0) (59) to perform a trajectory analysis of CD4 and CD8 T cell subsets using the slingshot (extend = n, stretch = 0, start.clus = Naïve) function on the UMAP embedding. Next Generalized Additive Models (GAMs) were employed to analyze gene expression dynamics along pseudotime in single-cell RNA sequencing (scRNAseq) data using the RunDynamicFeatures (n_candidates = 2000, seed = 2024, minfreq = 0) and DynamicHeatmap (use_fitted = TRUE, r.sq = 0.09, dev.expl = 0.1, num_intersections = NULL) functions from the SCP (v0.5.6) package (60). In brief, for each gene, a GAM was fit with gene expression as the response variable and pseudotime as the predictor, incorporating smooth functions to capture non-linear relationships. This approach enabled the identification of pseudotemporal changes in gene expression.

#### Myeloid Cells

To analyze myeloid cells and their various subsets, we extracted all cells labeled as “Mono/Mac” using the previously assigned broad labels. Given the large population size Neutrophils were excluded from the myeloid subset analysis and were analyzed separately. We largely followed the same approach used for the initial broad analysis with a few modifications. In brief, we identified 2000 HVGs. For dimension reduction, integration, and clustering we used 20 PCs and a cluster resolution of 0.5.

We annotated cells according to the methods previously outlined, utilizing marker genes, differential expression, and LungMAP resources. For trajectory analysis again we used the slingshot (extend = n, stretch = 0) function on the UMAP embedding. Lastly we generated a heatmap of the dynamic features along the pseudotime trajectory using functions RunDynamicFeatures (n_candidates = 2000, seed = 2024, minfreq=0), DynamicHeatmap (use_fitted = TRUE, n_split = 5, r.sq = 0.09, dev.expl = 0.1, num_intersections = NULL) from the SCP (v 0.5.6) package (60).

#### Neutrophils

To further analyze neutrophils and their various subsets, we followed the same approach used for the entire dataset with a few modifications. Given the smaller size of the Neutrophil subset we identified 2000 HVGs. For dimension reduction, integration, and clustering we used 10 PCs and a cluster resolution of 0.5. Following our earlier approach, cell annotation involved the use of marker genes, differential expression, and the LungMAP dataset. To perform trajectory analysis, we once again utilized the slingshot (extend = n, stretch = 0, start.clus = Circulating Neu.) function on the UMAP embedding. Lastly we generated a heatmap of the dynamic features along the pseudotime trajectory using functions RunDynamicFeatures (n_candidates = 2000, seed = 2024, minfreq=0), DynamicHeatmap (use_fitted = TRUE, n_split = 5, r.sq = 0.09, dev.expl = 0.1, num_intersections = NULL) from the SCP (v 0.5.6) package.

#### Relative Abundance Calculation

For each subset of the dataset, the total count of all cells was used to determine the final relative proportions. Specifically, the relative abundance of each cell type within each sample was calculated by grouping cells based on cell type and timepoint, determining cell counts, and computing the proportion of each cell type relative to the total cell count per sample. All proportions were calculated relative to the entire dataset. Visualization was performed using ggplot2.

#### Pathway and Functional Enrichment Analysis

Enrichment analysis was conducted using the Escape (v 1.99.1) (61) and fgsea (v 1.30.0) (62) packages. We curated relevant pathways from MSigDB, including KEGG, Reactome, Hallmark, and WikiPathways(63–68). For visualization purposes, we abbreviated the pathway sources as follows: KEGG as ’K’, Reactome as ’R’, Hallmark as ’H’, and WikiPathways as ’WP’ in all pathway figures. For the analysis, the runEscape (method = AUCell) function was employed utilizing the AUCell enrichment scoring approach (69). This method was selected for its speed and superior performance with single-cell data compared to traditional methods (70). For the enrichment analysis of non-immune cells, we conducted pairwise differential expression comparisons between each timepoint and the D0 timepoint using the FindMarkers function. Following the differential expression analysis, genes were ranked by multiplying the -log10 p-value by the sign of the log2 fold change. The ranked genes were then analyzed using the fgseaMultilevel function (minSize = 15, maxSize = 500, nproc = 1, nPermSimple = 10000) to perform pathway enrichment on selected pathways. Data visualization was carried out using the heatmapEnrichment function and ggplot for the fgsea enrichment analysis.

#### Cell Communication Analysis

Cell communication analysis was conducted using CellChat (v2.1.2) (71). Each time point was processed independently and later merged into a unified CellChat object for comprehensive analysis. The processing followed the standard suggested workflow with largely default parameters. Initially, the data was preprocessed for cell communication analysis by subsetting the dataset and identifying overexpressed genes and interactions using the subsetData, identifyOverExpressedGenes, and identifyOverExpressedInteractions functions. Communication probability was calculated using computeCommunProb, considering the effect of cell proportions (population.size = TRUE). Interactions involving cells with fewer than 10 cells were filtered out using filterCommunication. Cell communication was inferred at the pathway level, and the communication was aggregated using computeCommunProbPathway and aggregateNet. To identify incoming and outgoing signaling strengths, network centrality scores were computed, which allowed us to identify senders and receivers using netAnalysis_computeCentrality. Subsequently, incoming and outgoing signaling strength plots for each time point were generated using netAnalysis_signalingRole_scatter. These plots were used to extract the data frame containing the signaling roles for each cell type. The resulting data frames were then merged into a single data frame and plotted using ggplot2. To observe changes in signaling pathways over time, we utilized rankNet to generate a stacked bar plot displaying relative signaling strength. Finally, we used netVisual_aggregate to generate circle plots for select pathways.

### Spatial Library Preparation and Sequencing

Spatial transcriptomic analysis of non-infected (0 dpi) and infected (14 dpi) lungs was conducted using 10x Visium technology (10x Genomics). Lung sections, 5 µm thick, were mounted onto charged slides, deparaffinized, and stained with hematoxylin and eosin. The sections were imaged at 4x magnification using an ECHO Revolve microscope, and the tiles were stitched together using Affinity Photo (Serif). Subsequently, the lung sections were destained and decrosslinked to release RNA sequestered by the formalin fixation. The sections were then incubated with mouse-specific probes targeting the whole transcriptome (PN-1000365, 10x Genomics), allowing for the hybridization and ligation of each probe pair. Gene expression probes were released from the tissue and captured by spatially barcoded oligonucleotides on the Visium slide surface using the Visium CytAssist instrument. Gene expression libraries were prepared from each tissue section and sequenced using the Illumina NextSeq 2000. Data processing, including alignment, barcode assignment, and gene expression quantification, was performed using the 10x Genomics Space Ranger (v2.1.1) pipeline. Alignments were carried out using the reference genome package refdata-gex-mm10-2020-A for GRCm38 (mm10), provided by 10x Genomics. Additionally, the "Visium_Mouse_Transcriptome_Probe_Set_v1.0_mm10-2020-A.csv" probe set from 10x Genomics was utilized.

### Spatial Transcriptomics Analysis

#### Data processing and Analysis

Analysis of the spatial transcriptomic data was performed using Seurat. For QC, we filtered out spots containing fewer than 200 genes. Data was then normalized using version 2 of the SCTransform function (seed.use = 2024, vst.flavor="v2") (72, 73). For dimension reduction and clustering, we selected the first 20 PCs for downstream analysis. We leveraged our labeled scRNAseq data to perform spatial deconvolution. For this purpose, we used CARD, a method known for its accuracy, speed, and user-friendliness and applied it to both samples (74, 75). First, we created a CARD object using createCARDObject and performed the deconvolution using CARD_deconvolution all with default parameters. To visualize expression of certain genes we plotted them for both samples using Seurat’s SpatialFeaturePlot (image.alpha = 0.4, pt.size.factor = 2.5) function. Using some curated pathways as described above, we performed an gene set co-regulation analysis (GESECA) using fgsea. We first applied SCT transformation using SCTransform on 10,000 genes and then reduced the dimensions by performing a reverse PCA to capture the linear combinations of the cells (62). This was done using RunPCA with the following parameters: assay = "SCT", rev.pca = TRUE, npcs = 50, reduction.name = "pca.rev", and reduction.key = "PCR_". The pathway enrichment analysis was performed using geseca (minSize = 15, maxSize = 500, center = FALSE) function from fgsea. Visualization of the enriched pathways was performed using plotCoregulationProfileSpatial from fgsea.

### Statistical analyses

Data were analyzed and graphed using Prism (GraphPad, La Jolla, CA) software. For comparison of cell counts and flow cytometry analyses, significance was assessed using one-way ANOVA with Tukey’s multiple comparison test. An adjusted p value of 0.05 or less was considered significant, and significant differences are indicated as follows: *p < 0.05, ***p < 0.001.

## Data availability

The single cell RNA-sequencing and spatial data have been deposited at the NCBI Gene Expression Omnibus (GEO). We will provide GEO accession numbers upon acceptance.

## Acknowledgements

Funding for this research was provided by internal Lawrence Livermore National Laboratory Directed Research and Development funds (22-ERD-010 to D.R.W. and G.G.L.). The funders had no role in study design, data collection and analysis, decision to publish, or preparation of the manuscript. This work was performed under the auspices of the U.S. Department of Energy by Lawrence Livermore National Security, LLC, Lawrence Livermore National Laboratory under Contract DE-AC52-07NA27344

## Author contributions

O.A.D: Conceptualization, Methodology, Formal analysis, Visualization, Investigation, Data Curation, Writing - Original Draft

D.R.W: Conceptualization, Methodology, Resources, Investigation, Project administration, Writing - Original Draft, Funding acquisition, Supervision

G.G.L: Conceptualization, Methodology, Funding acquisition

K.K.H: Conceptualization

A.S: Conceptualization, Methodology, Formal analysis, Visualization, Data Curation

N.R.H: Conceptualization, Methodology, Investigation

N.F.L: Investigation, Visualization

N.M: Investigation,

M.V.R: Investigation, Visualization

A.M.P: Investigation, Data Curation

D.K.M: Investigation, Data Curation

All authors were involved in manuscript review and editing.

## Abbreviations

AM: Alveolar Macrophage
ANGPTL: Angiopoietin-Like Protein Family
C Mono: Classical Monocyte
C. posadasii: Coccidioides posadasii
CD4: Cluster of Differentiation 4
CD8: Cluster of Differentiation 8
Cd274: CD274 Antigen (PD-L1)
Cd5l+ Mac: CD5 Antigen-Like Macrophage
Circulating Neu: Circulating Neutrophils
COVID-19: Coronavirus Disease 2019
CXCL: Chemokine (C-X-C motif) Ligand
Cxcl2: Chemokine (C-X-C motif) Ligand 2
DCs: Dendritic Cells
dpi: days post infection
Early Neu A: Early Neutrophils A
Early Neu B: Early Neutrophils B
GESECA: Gene Set Co-Regulation Analysis
HVGs: Highly variable genes
IL-6: Interleukin 6
ILCs: Innate Lymphoid Cells
ISG: Interferon-Stimulated Gene
ISG Mono: Interferon-Stimulated Gene-expressing Monocyte
ISG Neu: ISG-expressing Neutrophils
Lyve1+ IM: Lymphatic Vessel Endothelial Hyaluronan Receptor 1 Interstitial Macrophage
MHCII IM: Major Histocompatibility Complex Class II Interstitial Macrophage
migDC: Migratory Dendritic Cell
Mono/Mac: Monocytes and Macrophages
NC Mono: Non-Classical Monocyte
NETosis: Neutrophil Extracellular Trap Formation
NK: Natural Killer
NKT: Natural Killer T cells
OPN: Osteopontin (Spp1)
OSM: Oncostatin M
OXPHOS: Oxidative Phosphorylation
PC: Principal Component
PCA: Principal Component Analysis
PD-L1: Programmed Death-Ligand 1
PD-L1+ Neu: PD-L1+ Neutrophils
Prolif.: Proliferating cells
S100a8: S100 Calcium Binding Protein A8
S100a9: S100 Calcium Binding Protein A9
scRNAseq: single-cell RNA sequencing
Spp1: Secreted Phosphoprotein 1 (OPN)
T/NK: T and Natural killer cells
Th1: T helper 1
Th17: T helper 17
TNF: Tumor Necrosis Factor
TNFA: Tumor Necrosis Factor Alpha
Tregs: Regulatory T cells
UMAP: Uniform Manifold Approximation and Projection

---

**Figure S1:**
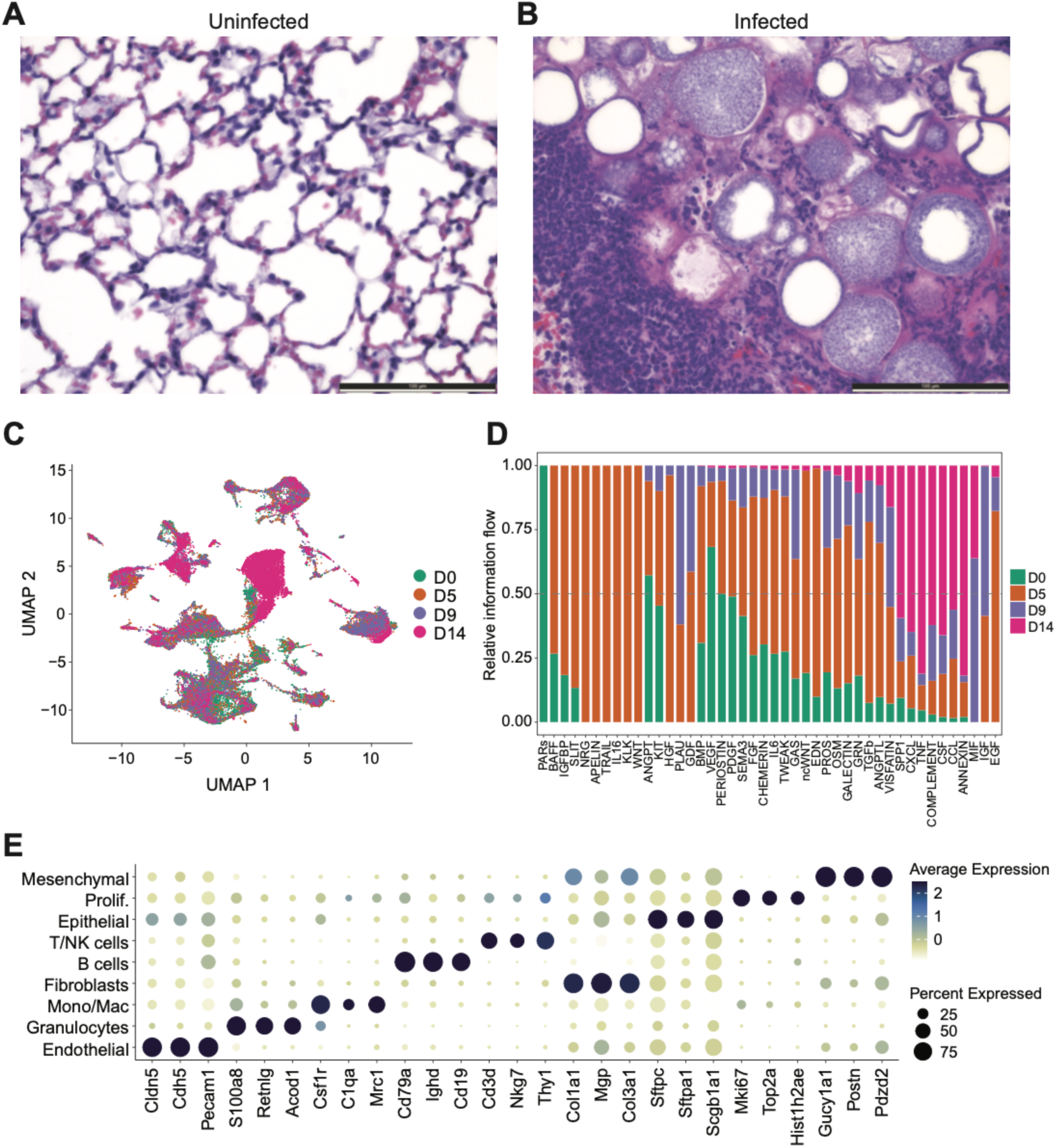
Supporting information for Figures 1 and 2. H&E staining of an uninfected (A) or infected (B) lung. Images taken at 40x magnification. The white scale bar in the bottom right-hand corner represents 100 µm. (C) UMAP plot showing the integration of multiple time points in the scRNAseq data set from lung samples. Cells are color-coded by time points: uninfected (0 dpi) and infected (5, 9, 14 dpi). (D) Stacked bar plot illustrating the relative information flow over time, defined as the total communication probabilities among cell types in the inferred network. The X-axis denotes the signaling pathways, while the Y-axis represents the relative information flow. Colors correspond to the different time points. (E) Dot plot illustrating the average expression levels of select genes. Expression levels are color-coded from low (muted yellow) to high (deep indigo). The size of each circle represents the percentage of cells expressing the gene.

**Figure S2:**
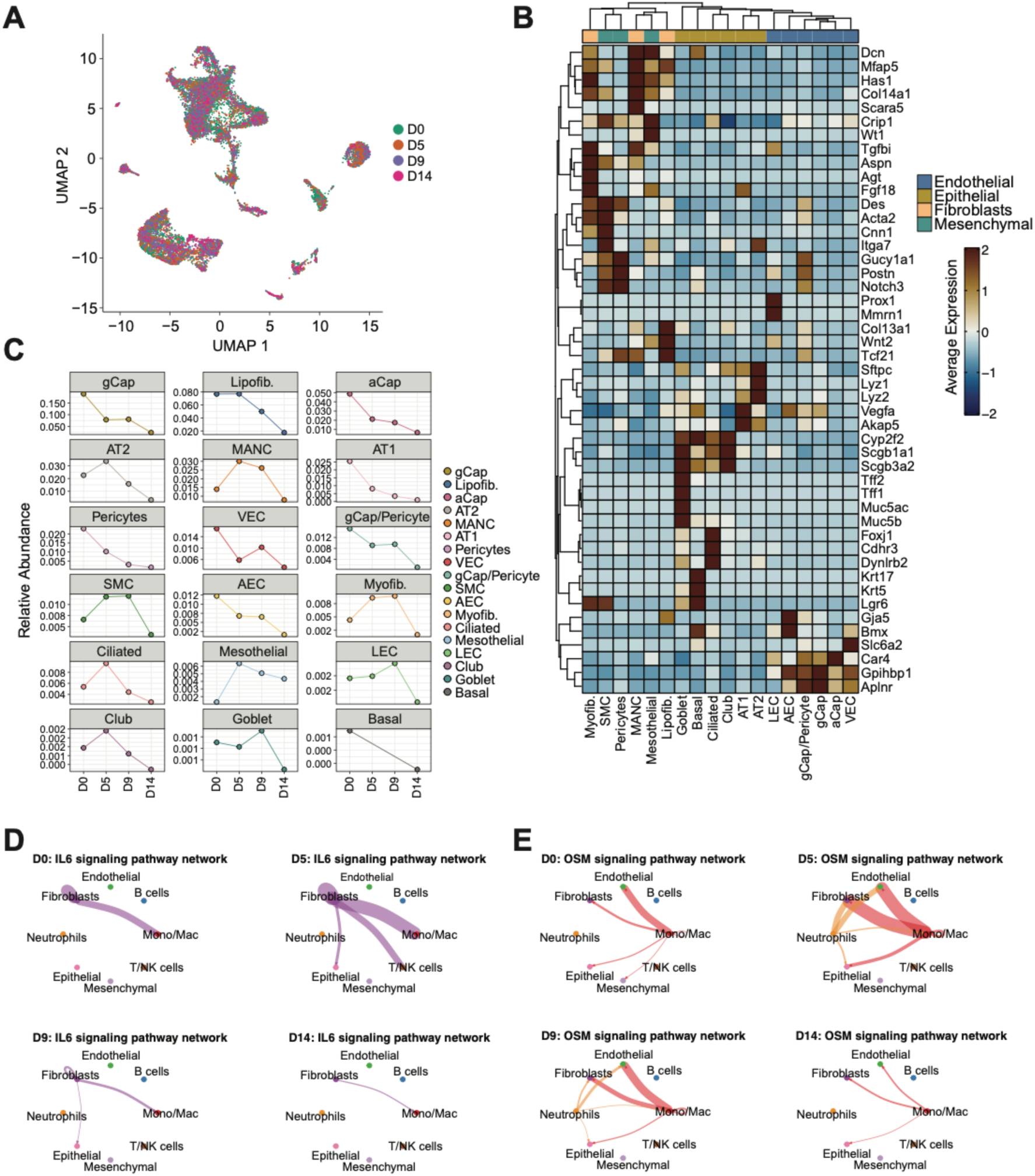
Gene expression and cell signaling pathways in non-immune cells. (A) UMAP plot of the non-immune cells showing the integration of multiple time points in the scRNAseq data set from lung samples. Cells are color-coded by time points: uninfected (0 dpi) and infected (5, 9, 14 dpi). (B) Heatmap illustrating the scaled average expression levels of selected genes. Expression levels are color-coded from low (dark blue) to high (dark brown). Column names represent the subtypes of the main non-immune cell types, with colors on top of each column indicating their respective groups. Row names correspond to the selected genes. Both genes and subtypes are hierarchically clustered based on expression similarity. (C) Faceted line plot depicting the relative abundance of each cell type in relation to all cells. Each subplot corresponds to a specific cell type and is color-coded accordingly. The X-axis represents the timepoints, while the Y-axis indicates the relative abundance. (D) Circle plots of IL6 signaling pathway network, with separate plots for each time point. Colors correspond to the main cell types. In each plot, edge colors represent the sources as the sender, and edge weights correspond to the interaction strength, with thicker lines indicating a stronger signal. (E) Circle plots of OSM signaling pathway network, with separate plots for each time point. Colors correspond to the main cell types. In each plot, edge colors represent the sources as the sender, and edge weights correspond to the interaction strength, with thicker lines indicating a stronger signal.

**Figure S3:**
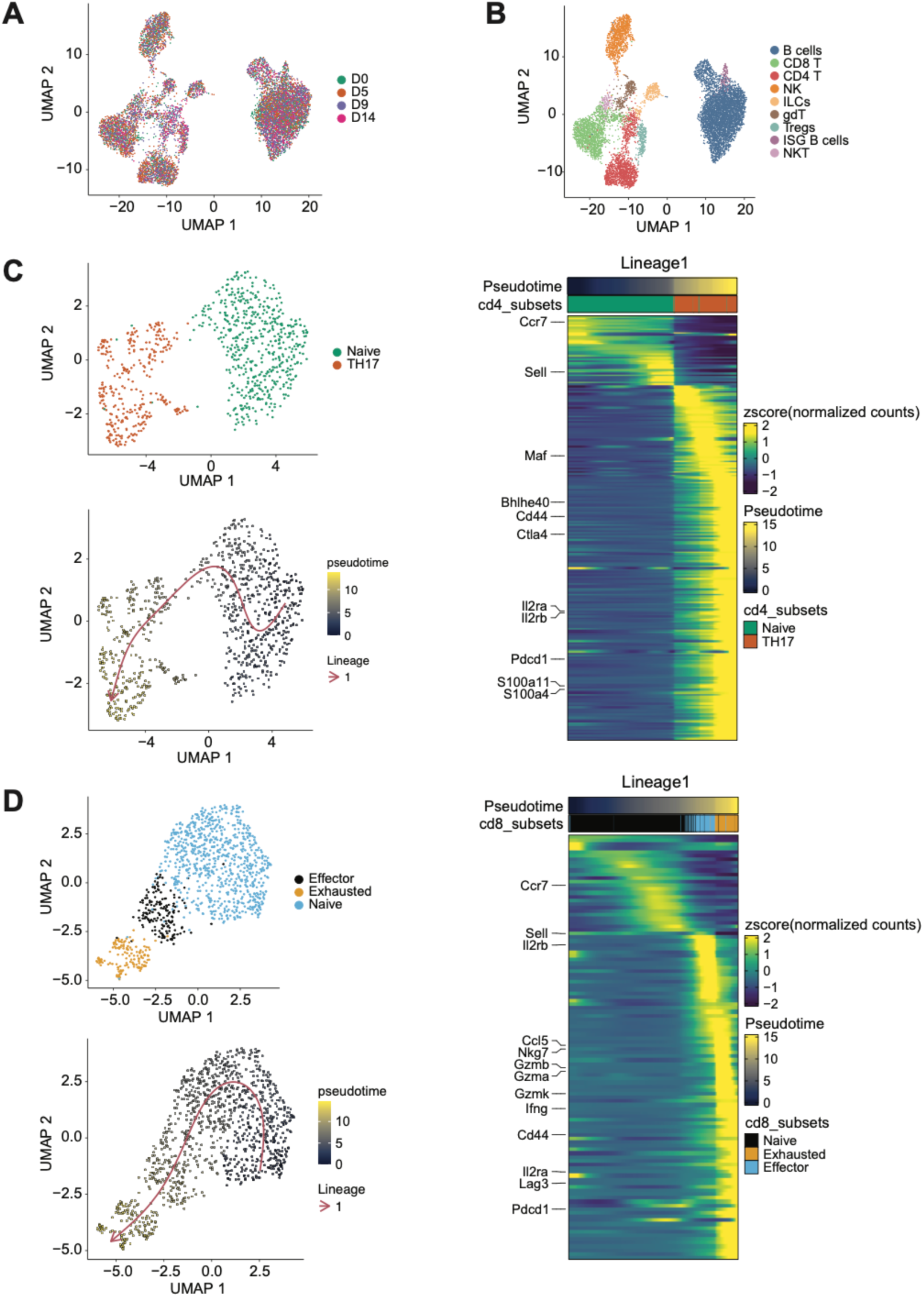
Lymphocyte trajectories and gene expression information. Lymphocyte subset integrated UMAP plots showing the integration of multiple time points in the scRNAseq dataset from lung samples. Cells are color-coded by time points (A) or cell type (B). CD4 (C) or CD8 (D) T cell subset UMAP plots displaying subtypes (top) and pseudotime trajectory (bottom). Cells are color-coded based on their pseudotime values, ranging from blue (early) to yellow (late). A heatmap illustrates the scaled fitted gene expression values along pseudotime (right), with expression levels color-coded from low (purple) to high (yellow). The top row panel represents pseudotime, ordered from left to right and color-coded based on pseudotime values, ranging from blue (early) to yellow (late). The next row shows cell type colors, where each bar represents a cell belonging to a specific cell type along the pseudotime trajectory. Rows represent genes, with selected genes highlighted on the left-hand side.

**Figure S4:**
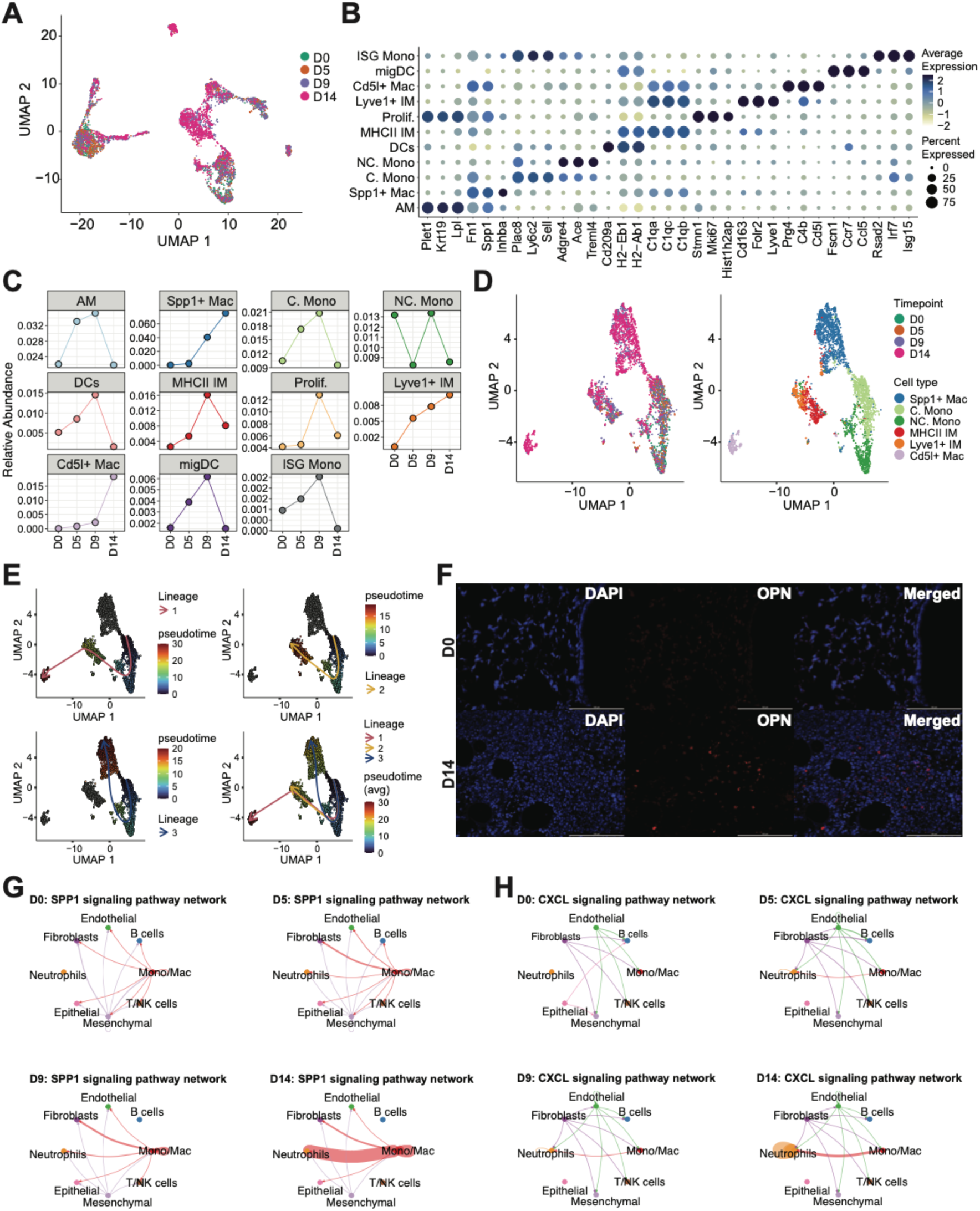
Gene expression, pseudotime trajectories, and cell signaling within myeloid cell populations. (A) Mono/Mac subset UMAP plot showing the integration of multiple time points in the scRNAseq data set from lung samples. Cells are color-coded by time points. (B) Dot plot illustrating the average expression levels of select genes. Expression levels are color-coded from low (muted yellow) to high (deep indigo). The size of each circle represents the percentage of cells expressing the gene. (C) Faceted line plot depicting the relative abundance of each cell type in relation to all cells. Each subplot corresponds to a specific cell type and is color-coded accordingly. The X-axis represents the timepoints, while the Y-axis indicates the relative abundance. (D) UMAP plots of select Mono/Mac cell types showing the integration of multiple time points (left) and the cell types (right). Cells are color-coded by time points or by cell type. (E) UMAP plots displaying pseudotime trajectories. Cells are color-coded based on their pseudotime values, ranging from blue (early) to red (late). The pseudotime trajectory was inferred in an unsupervised manner using Slingshot. Each arrow represents a distinct identified trajectory (lineage). (F) IHC of the lungs at 0 dpi and 14 dpi. DAPI staining is represented in blue, OPN (*Spp1*) is represented in red, and the merged images show both colors. The white scale bar in the bottom right-hand corner represents 100 µm. Images taken at 40x magnification. Circle plots of SPP1 (G) or CXCL (H) signaling pathway network, with separate plots for each time point. Colors correspond to the main cell types. In each plot, edge colors represent the sources as the sender, and edge weights correspond to the interaction strength, with thicker lines indicating a stronger signal.

**Figure S5:**
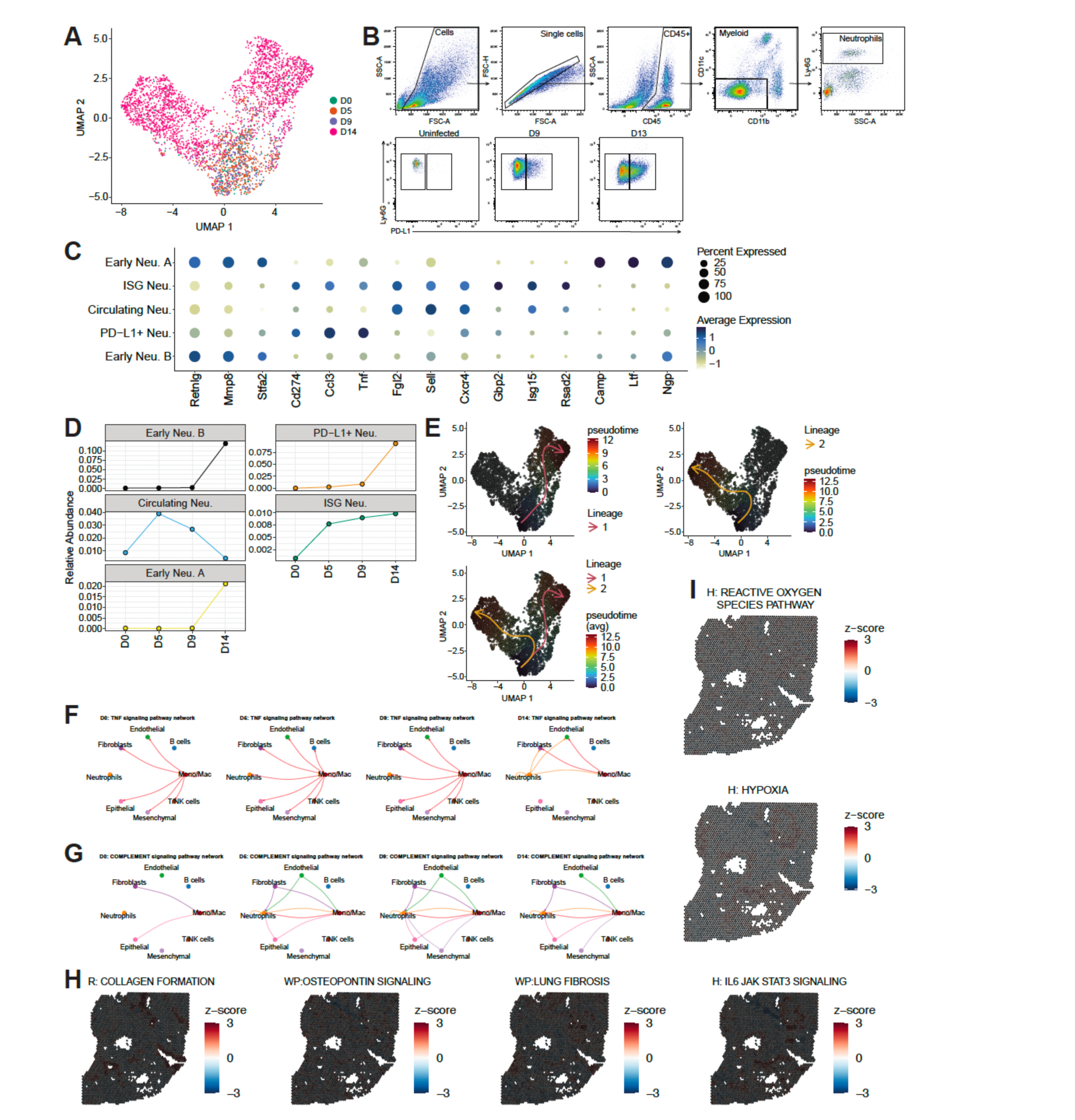
Gene expression and pseudotime trajectories of neutrophils, and supplemental spatial sequencing data. (A) An integrated UMAP plot of subclustered neutrophil sub-populations in the lung from uninfected (0 dpi) and infected (5, 9, 14 dpi) samples. Cells are color-coded by time point. (B) Gating strategy for the identification of PD-L1^+^ neutrophils. (C) Dot plot illustrating the average expression levels of select genes. Expression levels are color-coded from low (muted yellow) to high (deep indigo). The size of each circle represents the percentage of cells expressing the gene. (D) Faceted line plot depicting the relative abundance of each cell type in relation to all cells. Each subplot corresponds to a specific cell type and is color-coded accordingly. The X-axis represents the timepoints, while the Y-axis indicates the relative abundance. (E) UMAP plots displaying pseudotime trajectories. Cells are color-coded based on their pseudotime values, ranging from blue (early) to red (late). The pseudotime trajectory was inferred in an unsupervised manner using Slingshot. Each arrow represents a distinct identified trajectory (lineage). Circle plots of TNF (F) or COMPLMENT (G) signaling pathway network, with separate plots for each time point. Colors correspond to the main cell types. In each plot, edge colors represent the sources as the sender, and edge weights correspond to the interaction strength, with thicker lines indicating a stronger signal. (H) and (I) Spatial plots of gene set co-regulation analysis (GESECA) on 14 dpi lung for selected pathways. A high positive z-score (red) indicates that the gene set is significantly enriched, while a low or negative z-score (blue) suggests that the gene set is not significantly enriched or is even depleted. Pathways from MSigDB (KEGG, Reactome, Hallmark, and WikiPathways) were curated and abbreviated as K, R, H, and WP, respectively.

